# The ULK1 effector BAG2 regulates autophagy initiation by modulating AMBRA1 localization

**DOI:** 10.1101/2023.12.08.570815

**Authors:** Devanarayanan Siva Sankar, Stephanie Kaeser-Pebernard, Christine Vionnet, Sebastian Favre, Lais Oliveira De Marchioro, Benjamin Pillet, Jianwen Zhou, Werner Josef Kovacs, Dieter Kressler, Manuela Antonioli, Gian Maria Fimia, Jӧrn Dengjel

**Author notes:** Corresponding author: Jörn Dengjel.

## Abstract

Canonical autophagy is regulated by ULK1, the most proximal protein kinase specifically regulating autophagy initiation. To gain new insights into functions of the ULK1 holo-complex in autophagy regulation, we generated a deep ULK1 complex interactome by combining affinity purification- and proximity labelling-mass spectrometry of all four ULK1 complex members: ULK1, ATG13, ATG101 and RB1CC1/FIP200. Under starvation conditions, the ULK1 complex interacts with several protein and lipid kinases and phosphatases implying the formation of a signalosome. Interestingly, also several selective autophagy receptors interact with ULK1 indicating the activation of selective autophagy pathways by nutrient starvation. One effector of the ULK1 complex is the HSC/HSP70 co-chaperone BAG2, which regulates the subcellular localization of the VPS34 lipid kinase complex member AMBRA1. Depending on the nutritional status, BAG2 has opposing roles. In growth promoting conditions, the unphosphorylated form of BAG2 sequesters AMBRA1, attenuating autophagy induction. In starvation conditions, ULK1 phosphorylates BAG2 on Ser31, supporting its recruitment to the ER membrane and positively affecting autophagy flux.

## Introduction

Macroautophagy (hereafter referred to as autophagy) is an evolutionary-conserved lysosomal degradation pathway critical for cellular homeostasis (Yamamoto et al., 2023). Autophagy is a constitutive process as well as a stress response which mainly fulfils cytoprotective functions. Its dysregulation is often observed in diseases such as cancer and neurodegeneration (Klionsky et al., 2021). A hallmark of autophagy is the *de novo* formation of double membrane vesicles, autophagosomes, which enwrap cytoplasm destined for degradation. Autophagosome biogenesis can be classified by five phases: initiation, membrane nucleation, expansion, pore closure/maturation, and fusion with lysosomes (Hu and Reggiori, 2022). Each phase is characterized by the involvement of specific genes and respective proteins which are regulated on transcriptional, translational, as well as posttranslational level (Delorme-Axford and Klionsky, 2018; Licheva et al., 2022). Immediately after a given stress stimulus, autophagosome initiation does not require *de novo* protein synthesis and is mainly regulated on a posttranslational level by a set of protein- and lipid-kinases (Abeliovich et al., 2000). The serine/threonine kinase unc-51 like autophagy activating kinase 1 (ULK1, or its homolog ULK2), the mammalian homolog of yeast autophagy-regulated (Atg) 1 (Chan et al., 2007; Young et al., 2006), is the most proximal protein kinase specifically regulating canonical autophagosome biogenesis. The catalytic subunit ULK1 along with ATG13 (Chan et al., 2009; Hosokawa et al., 2009a), and ATG101 (Hosokawa et al., 2009b; Mercer et al., 2009) forms an asymmetric complex with RB1CC1/FIP200 (Hara et al., 2008), with a 1:1:1:2 stoichiometry (Shi et al., 2020). Upon autophagy induction, this complex is recruited to ER subdomains (Karanasios et al., 2016), where it phosphorylates several proteins important for functional autophagy, amongst others members of the lipid kinase VPS34 complex I (Mercer et al., 2018; Mercer et al., 2021).

Due to technical and methodological improvements, mass spectrometry (MS)-based proteomic approaches continue to generate new insights into ULK1-dependent signal transduction (Hu et al., 2021; Mercer et al., 2021). In the current study, we decided to employ state-of-the-art MS-based proteomics to revisit ULK1 complex protein-protein interactions. In contrast to the majority of published approaches which used classical affinity purification (AP)-MS (Behrends et al., 2010; Joo et al., 2016), we used a combination of AP- and proximity labeling (PL)-MS to generate a deep interactome of the ULK1 holo-complex. PL-MS relies on the expression of proteins-of-interest fused to enzymes covalently transferring chemical groups to proximal proteins labeling so called protein neighborhoods. This supports the identification of weak transient interactions in a radius of 10-20 nm around bait proteins (for review see (Siva Sankar and Dengjel, 2021)). Using this double strategy, we generated a comprehensive picture of ULK1 complex interactions characterizing common and subunit-specific interaction partners. Corroborating data from signaling studies (Hu et al., 2021; Mercer et al., 2021), the function of ULK1 appears to be more complex, not only affecting autophagosome initiation, but also later phases of the organelle life cycle.

One protein that caught our attention and that was studied in detail is Bcl-2-associated athanogene 2 (BAG2), which interacts via its BAG domain with HSC/HSP70 proteins, acting as a nucleotide exchange factor (Takayama and Reed, 2001). BAG2 is a member of the evolutionary conserved BAG family, which in humans consists of six members BAG1-6. BAG proteins are suggested to act antiapoptotic and were shown to modulate proteasomal as well as autophagosomal protein degradation (Kabbage and Dickman, 2008; Pattingre and Turtoi, 2022). BAG2 is widely expressed (Munthe et al., 1996) and localizes to various subcellular locations such as mitochondria (Qu et al., 2015), ER (Arndt et al., 2005), and microtubules (de Paula et al., 2016). It seems to modulate the balance between chaperone-mediated protein folding and chaperone-mediated protein degradation, amongst others by inhibiting CHIP, an HSC/HSP70 chaperone-associated ubiquitin ligase (Arndt et al., 2005). Similar as BAG3 and BAG6, BAG2 was shown to promote mitophagy (Che et al., 2013; Qu et al., 2015; Ragimbeau et al., 2021; Tahrir et al., 2017) and reticulophagy via its interaction with the selective autophagy receptor (SAR) SQSTM1/p62 in the context of *Mycobacterium tuberculosis* infection (Liang et al., 2020).

In the current manuscript, we characterize BAG2 as ULK1 complex interactor, target and downstream effector, one of its autophagy-relevant clients being the VPS34- and E3 ubiquitin ligase complex-member Activating molecule in BECN1-regulated autophagy protein 1 (AMBRA1) (Antonioli et al., 2017). Under growth conditions BAG2 sequesters AMBRA1 attenuating autophagy; however, upon autophagy activation, BAG2 translocates to ER subdomains supporting functional autophagy. Changes in the intracellular localization of BAG2 are regulated by ULK1 via phosphorylation of BAG2 at Ser31.

## Results

### The ULK1 holo-complex interacts with selective autophagy receptors in starvation conditions

Protein-protein interactions of ULK1 complex members have been analyzed by AP- and PL-MS in low and high throughput studies (Behrends et al., 2010; Tu et al., 2021; Wang et al., 2019) (for review see (Siva Sankar and Dengjel, 2021)). Depending on the exact experimental setup (e.g., AP of endogenous or tagged-proteins, tag localization), different sets of interaction partners might be identified. To generate a comprehensive overview of ULK1 complex interactions in one cell line under one physiological condition i.e., nutrient starvation, we decided to generate a deep ULK1 complex interactome by combining AP- and PL-MS of N- and C-terminal tagged fusion proteins. Firstly, using CRISPR/Cas9 we generated four HeLa knockout (KO) cell lines, one for each complex member i.e., ULK1, ATG13, ATG101 and FIP200, respectively (Figure 1A). As anticipated, all four cell lines exhibited a block in autophagy as analyzed by accumulation of p62/SQSTM1 under starvation conditions (Figure 1B). Interestingly, loss of single complex members had differential effects on abundances of the other members (Figure 1A): loss of FIP200 led to a downregulation of all the other complex members, supporting its central function in complex assembly (Shi et al., 2020). In contrast, loss of ATG101 and ULK1 led to compensatory effects and increased abundances of the other members. Loss of ATG13 had a positive effect on the abundance of FIP200 and negative effects on ULK1 and ATG101 abundances (Goodwin et al., 2017; Hosokawa et al., 2009b). These data imply that the holo-complex forms by preassembled subcomplexes (Shi et al., 2020). Loss of complex members also affected ULK1 activity differentially as analyzed by phosphosite-specific western blots against the ULK1 target site on ATG14, Ser29 (Figure 1A, pATG14). Whereas knockout of ULK1, ATG13 and FIP200 led to reduced ATG14 phosphorylation, knockout of ATG101 did not. This implies that loss of ATG101 does not interfere with ULK1 complex assembly and activity per se, but rather negatively interferes with the recruitment of downstream components such as ATG9 (Nguyen et al., 2023), which ultimately also leads to a block of functional autophagy (Figure 1B).

**Figure 1:**
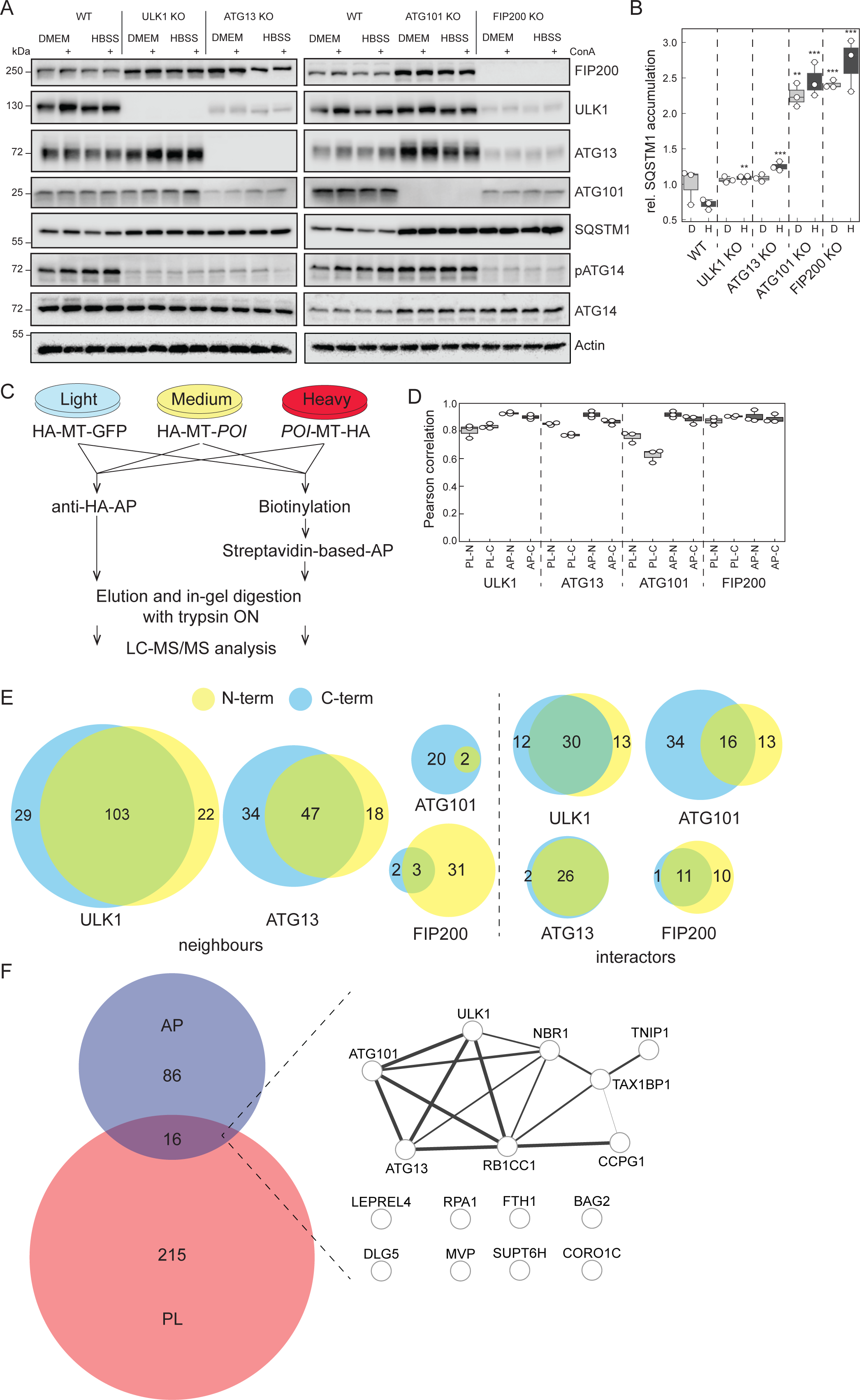
Deep interactome of the ULK1 holo-complex. **(A)** Generation of HeLa KO cell lines of ULK1 complex members. Using CRISPR/Cas9 and specific single guide RNAs (sgRNAs), KO cells for ULK1, ATG13, ATG101 and RB1CC1/FIP200 (proteins-of-interests [POIs]) were generated, respectively. Shown is one representative western blot of n=3 biological replicates. ConA: concanamycin A. pATG14 = phosphorylated Ser29 of ATG14. **(B)** Blocked autophagy in KO cell lines. Quantification of blots shown in (A) normalized to actin. All KO cells exhibit an accumulation of SQSTM1/p62 in starvation conditions indicative of blocked autophagy. Box plots show spreads of relative protein abundances. Single values are highlighted as white dots. D: DMEM; H: HBSS; n=3 biol. replicates, Student’s t-test, unpaired, **: p<0.01; ***: p<0.001. **(C)** Quantitative proteomics workflow. KO cells were stably transfected with doxycyclin (dox)-inducible expression constructs encoding N- and C-terminal tagged fusion POIs, employing a combination of the hemagglutinin (HA) and miniTurbo (MT) tags. A SILAC-based approach was used to generate deep interactomes comparing in single MS experiments cells expressing N- or C-terminal tagged POIs to cells expressing a MT-tagged GFP variant as negative control. Anti-HA affinity purifications (AP) as well as MT-based biotinylation reactions of proximal proteins (proximity labeling [PL]) followed by streptavidin-based AP were performed prior to quantitative LC-MS/MS analyses. **(D)** Correlation of biological replicates. For each POI and each tag-variant n=3 biological replicates were performed. Box plots show spreads of Pearson correlation coefficients. Single values are highlighted as white dots. **(E)** Proportional Venn diagrams highlighting overlap of significantly enriched neighbors and interactors of indicated experiments (p<0.05, see Methods for details). **(F)** Proportional Venn diagram highlighting overlap of significantly enriched interactors identified by AP-MS and enriched neighbors identified by PL-MS. The 16 commonly identified proteins were analyzed by STRING DB on known protein-protein interactions. Nodes represent proteins and edge thickness confidence of interactions.

In each KO cell line, we introduced an inducible expression system supporting the doxycyclin (dox)-controlled expression of either the respective N-terminal or C-terminal tagged protein-of-interest (POI) (supplemental Figure S1). As tag we used a combination of the peptide epitope of human influenza virus hemagglutinin (HA) and the biotin ligase variant miniTurbo (MT) (Branon et al., 2018), supporting both AP- and PL-analyses using a single setup (Liu et al., 2020) (Figure 1C). Using stable isotope labeling by amino acids in cell culture (SILAC)-based proteomics we determined the interactions of ULK1 complex members under starvation conditions (90 min of HBSS treatment), the classical stimulus for the induction of bulk autophagy, by comparing POI-to HA-MT-GFP-based enrichments as negative controls (Figure 1C). For each analysis i.e., AP- and PL-MS of N- and C-terminal tagged POI, we performed three biological replicates, which led to a total of 48 LC-MS/MS experiments, 24 AP- and 24 PL-MS studies (supplemental Tables S1A-B). Biological replicates correlated well with an average Pearson correlation coefficient of 0.85 (Figure 1D, range from 0.57-0.93). Interestingly, the PL-MS studies of ATG101 showed reproducibly the lowest correlation (Figure 1D, 0.57-0.78). ATG101 is with 25 kDa by far the smallest protein of the complex and the low correlation of transient interactions identified by PL-MS might indicate a high mobility of ATG101. High-confidence interactors were defined as proteins which were identified as significantly enriched compared to the MT-GFP negative control in minimally two AP- or two PL-MS experiments, respectively (Significance A, p<0.05, BH corrected for PL-MS, see Methods for details). As expected, PL-MS analyses yielded more significantly enriched neighboring proteins compared to interactors identified by AP-MS, one exception being ATG101 (Figure 1E). The site of the tag did have an influence on the identified interactions in both approaches. The most prominent differences were observed for the PL-MS analyses of ATG101 and FIP200 (Figure 1E). Whereas the N-terminal tagged version of FIP200 revealed 34 neighboring proteins, only five neighbors were enriched with the C-terminal variant. This reduced efficiency might be linked to the structure of the ULK1 holo-complex, in which two FIP200 protomers form a homodimer via their C-terminal regions and claw domains (Shi et al., 2020; Turco et al., 2019). The C-terminal tag might interfere with holo-complex formation. For ATG101 the C-terminal tagged variant yielded a higher number of potential neighbors, which might highlight a better accessibility compared to the N-terminal variant (Qi et al., 2015).

In total we identified 317 proteins as neighbors and interactors (Figure 1F, supplemental Table S1C-D). Compared to other published ULK1 complex interactomes we cover 65 of the known and characterize 252 new interactors (Supplemental Figure S2A) (Behrends et al., 2010; Tu et al., 2021; Wang et al., 2019), the overlap of this study to the published works being larger than the overlap between these. As has been described for AP- and PL-MS studies (Lambert et al., 2015; Liu et al., 2020), both approaches are highly complementary and uncover different sets of interacting and neighboring proteins. Using our stringent selection criteria only 16 proteins i.e., ca. 5% of the entire dataset, were identified by both approaches, amongst them the four holo-complex members themselves (Figure 1F). Intriguingly, the selective autophagy receptors (SARs) CCPG1, NBR1, and TAX1BP1 appear to interact closely with the ULK1 complex as they were among the 16 commonly enriched proteins (Kirkin et al., 2009; Newman et al., 2012; Smith et al., 2018). In selective autophagy, the recruitment of the ULK1 complex to degradation-primed cargo via interaction between FIP200 and SARs has been recognized as a decisive event (Ohnstad et al., 2020; Ravenhill et al., 2019; Turco et al., 2019; Zhou et al., 2021), TAX1BP1 being the main driver by FIP200 binding (Turco et al., 2021). Our finding indicates that also under starvation-induced autophagy i.e., the prototypical stimulus of non-selective bulk autophagy, selective removal of cargo might take place.

### The ULK1-complex forms a signaling hub

The best-known function of the ULK1 complex is the induction of autophagy. Other functions are less well understood; however, analyses of its molecular targets indicate that it regulates also later stages of autophagy, such as autophagosome-lysosome fusion (Hu et al., 2021; Sanchez-Martin et al., 2023), and that it may affect other pathways, ranging from glycolysis (Li et al., 2016) to mRNA splicing (Schmitz et al., 2021). Our deep interactome revealed that out of the 317 interactors, 102 interacted with minimally two complex members and 215 were member-specific interactors (Figure 2A). As a quality check we scanned the interactome for proteins with known functions in autophagy and were able to generate a subnetwork of 31 proteins i.e., ca. 10% of the entire dataset (Figure 2B). These proteins were mainly linked to early events in autophagy, such as membrane nucleation (PIK3C3, ATG16L1) and expansion (ATG2B, ATG5, WIPI2), and to cargo recruitment. Next to the SARs mentioned above we identified p62 (Sanchez-Martin et al., 2019), interestingly only by AP-MS and not by PL-MS, as well as NCOA4 and FTH1, which regulate ferritin degradation and iron homeostasis (Dowdle et al., 2014). This subnetwork highlights the successful enrichment of autophagy-relevant proteins and supports our experimental and data analysis pipeline with the aim of identifying new proteins being important in autophagy regulation.

**Figure 2:**
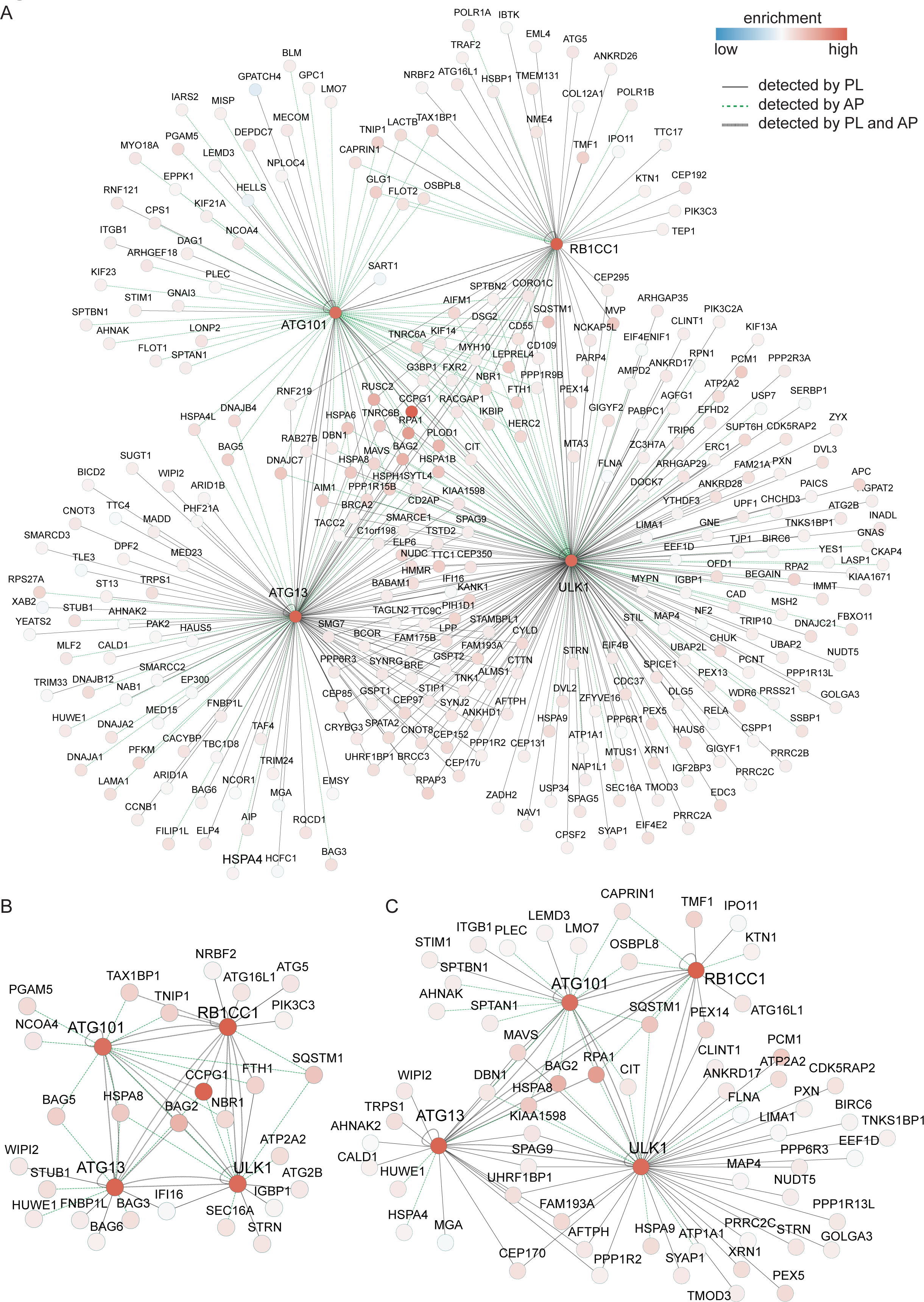
The ULK1 holo-complex forms a signaling hub. **(A)** Deep ULK1 complex interactome. Shown are significant protein-protein interactions identified by AP- and PL-MS under starvation conditions (90 min of HBSS treatment). The node color-code indicates the relative enrichment in all 48 experiments highlighted in Figure 1D. The edge color-code indicates the type of performed experiments (AP-MS, PL-MS, or both). **(B)** ULK1 complex interactors with known functions in autophagy. **(C)** ULK1 complex interactors that were identified as potential ULK1 targets carrying ULK1-regulated phosphosites.

The identified 317 proteins are linked to numerous subcellular localizations as well as to many functions, indicating a much broader role of ULK1 in shaping the cellular response to starvation. We would like to highlight some interesting observations using the presented data as a resource to generate new hypotheses: the wealth of interactions indicates that the ULK1 complex acts as a signaling hub i.e., forms a macromolecular signalosome. We identified the ULK1 complex as interacting with five protein-, two lipid-, and two metabolite-kinases (supplemental Figure S2B). Whereas the PI3P generating kinase PIK3C3/VPS34 is a well-known target of ULK1 (Egan et al., 2015), it is not known if ULK1 also phosphorylates PIK3C2A, another PI3P generating kinase which has been shown to induce autophagy in response to shear stress (Boukhalfa et al., 2020). The alpha subunit of the inhibitor of NFκB kinase CHUK/IKKA has been shown to be degraded by autophagy (Tan et al., 2020) and to promote AMBRA1-ATG8 interaction in mitophagy (Di Rita et al., 2018); however, a direct crosstalk with ULK1 has not been described. This is also true for the tyrosine protein kinase YES1, which was shown to promote autophagy in ovarian cancer (Zhou et al., 2022), and for the protein kinases CIT, PAK2 and TNK1, which have been linked amongst others to cytoskeleton regulation but not to autophagy.

The ULK1 complex also interacts with eleven regulatory phosphatase subunits (supplemental Figure S2C). Four of these, PPP1R2, PPP6R3, PPP1R13L and STRN were identified by us as ULK1 targets, the PP2a regulatory subunit STRN being involved in positive feedback promoting the dephosphorylation and activation of ULK1 (Hu et al., 2021). Finally, we scanned the entire interactome for known ULK1 targets and generated a subnetwork consisting of the four complex members and 60 published target proteins (Figure 2C) (Alsaadi et al., 2019; Egan et al., 2015; Hu et al., 2021; Joo et al., 2016; Li et al., 2016; Mercer et al., 2021). The network contains proteins related to autophagy (FDR=0.0129), such as p62, ATG16L1, WIPI2, but also many proteins related to cytoskeleton (FDR=0.00031) and intracellular transport (FDR=0.0071), indicating that ULK1 also regulates autophagosome trafficking. Taken together, we generated a comprehensive list of proteins interacting or being in close spatial proximity of the ULK1 holo-complex under starvation conditions. Many of these are ULK1 targets and can be related to autophagy regulation or signal transduction supporting the interpretation that ULK1 complexes form macromolecular signalosomes.

### The co-chaperone BAG2 is a ULK1 interactor which modulates functional autophagy

The HSC/HSP70 co-chaperone BAG2 caught our attention (Takayama et al., 1999), as it was one of the 16 proteins that were identified by both AP- and PL-MS approaches (Figure 1F) and as it carries a potential ULK1 phosphorylation site (Figure 2C). The family of BAG proteins is conserved from yeast to mammals and has important functions in protein homeostasis (Pattingre and Turtoi, 2022). BAG2 was shown to support p62-dependent reticulophagy in *Mycobacterium tuberculosis*-infected macrophages by interfering with the Beclin1/BCL2 interaction, releasing Beclin1 to support autophagy (Liang et al., 2020); however, its role in classical starvation-induced autophagy and the underlying molecular mechanisms are largely unknown.

As indicated above, we identified BAG2 as significant interaction partner of ULK1, ATG101 and ATG13 by MS analyses (Figure 2A), as well as by biotin-based AP-western blot (supplemental Figure S1). To test if these interactions are potentially direct, we expressed BAG2 and the three ULK1 complex members in yeast and performed two-hybrid assays. Only in the case of the C-terminal domain of ULK1 (amino acids 827-1050, abbreviated as ULK1.827C) we observed activation of the *HIS3* and *ADE2* reporter genes and, thus, growth on selective medium lacking histidine or adenine, indicating that the ULK1-BAG2 interaction is likely direct. The interactions of BAG2 with ATG13 and ATG101 might be indirect as we did not observe activation of the reporter genes (supplemental Figures S3). In ULK1 AP-MS experiments, BAG2 was enriched to a similar extent as the other complex members (Figure 3A). To confirm these findings, we tested the interactions in a reverse co-AP of ectopically expressed tagged BAG2 followed by western blot analysis against ULK1 and ATG13 (Figure 3B-C, n=3). Starvation appeared to stabilize the interaction of BAG2 and ULK1; however, the difference was not statistically significant. As the BAG2-ATG13 interaction was similar in growth and in starvation conditions, we conclude that BAG2 is likely a constitutive interaction partner of the ULK1 complex (Figure 3B-C).

**Figure 3:**
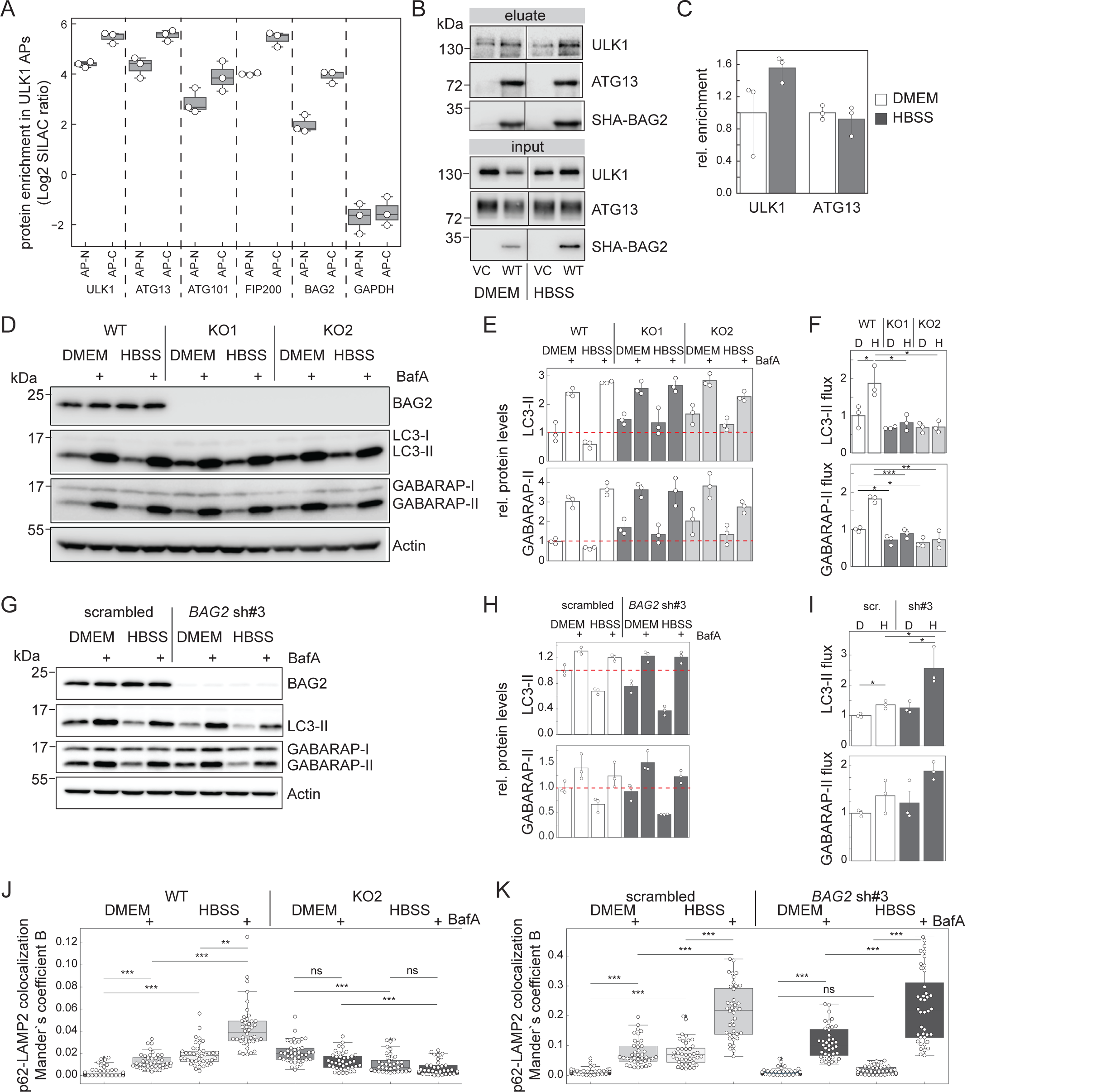
BAG2 is a constitutive interaction partner of ULK1 modulating autophagy. **(A)** BAG2 binds similarly to ULK1 as its complex members as identified by AP-MS. Shown are normalized and log2 transformed SILAC ratios of AP-MS experiments using N- and C-terminally tagged ULK1 variants. GAPDH is depicted as negative control (n=3 biological replicates each). Box plots show spreads of ratios. Single values are highlighted as white dots. **(B-C)** Co-IP of SHA-tagged BAG2 and its interaction partners ULK1 and ATG13. Shown is one representative experiment of n=3 biological replicates. Equal protein amounts per lane were loaded. Note: samples were run on one gel, black lines indicate cropping of unrelated lanes. (C) Quantification of n=3 replicates. Single values are highlighted as white dots. Error bars indicate 95% confidence interval. **(D-F)** *BAG2* KO reduces autophagy flux. Shown is one representative blot of n=3 biological replicates (D). Actin serves as loading control. Cells were starved in HBSS and/or treated with BafA1 for 3 h. Respective quantification of LC3-II and GABARAP-II bands are shown in (E). Red dotted lines indicate protein levels in DMEM control conditions. Comparing BafA1-treated with non-treated lanes indicates reduced autophagy flux in KO compared to WT cells (F). Student’s t-test, unpaired, *: p<0.05, **: p<0.01, ***: p<0.001. Single values are highlighted as white dots. Error bars indicate 95% confidence interval. **(G-I)** *BAG2* KD increases autophagy flux. Shown is one representative blot of n=3 biological replicates (G). Actin serves as loading control. Cells were starved in HBSS and/or treated with BafA1 for 3 h. Respective quantification of LC3-II and GABARAP-II bands are shown in (H). Red dotted lines indicate protein levels in DMEM control conditions. Comparing BafA1-treated with non-treated lanes indicates increased autophagy flux in KD compared to control cells (I). Student’s t-test, unpaired, *: p<0.05. Single values are highlighted as white dots. Error bars indicate 95% confidence interval. **(J-K)** Colocalization of p62 and LAMP2 analyzed by IF. In WT and KD cells colocalization increases upon starvation (HBSS, 3 h) and upon BafA1 treatment (2 nM, 3 h). This is not the case in KO cells. Data of n=3 biological replicates each, 20 cells per replicate. White dots represent the colocalization foci count per cell and per biological replicate. An unpaired two-tailed Student’s t-test was used to compare values. ns: not significant, *: p **≤** 0.05, **: p **≤** 0.01, ***: p **≤** 0.001. Respective fluorescence micrographs are shown in supplemental Figure S5.

We next asked whether BAG2 supports functional autophagy. For this we generated (i) CRISPR/Cas9-based HeLa KO cells using single guide RNAs (for sgRNA sequences see supplemental Table S2), and (ii) HeLa knockdown (KD) cells using a shRNA-based approach. With respect to KO cells, we generated two cell clones that were negative for BAG2 using limiting dilution. We first analyzed autophagy flux in KO cells monitoring MAP1LC3A/B (LC3) and GABARAP lipidation levels in the absence and presence of Bafilomycin A1 (BafA1), which blocks lysosomal acidification and thus inhibits lysosomal protein degradation (Figure 3D-F). Both clones exhibited an accumulation of lipidated LC3 and GABARAP in the absence of BafA1, especially under starvation conditions (Figure 3D-E). As the maximal lipidation level under BafA1 treatment did not increase in BAG2 KO clones compared to wildtype (WT) cells, this led to a reduction in autophagosome turnover, i.e., autophagy flux (Figure 3F). In KO cells, a decreased autophagy flux under starvation conditions was also observed by monitoring LC3 and p62 protein abundances by MS (supplemental Figure S4A), as well as by p62 puncta accumulation by immunofluorescence (IF) (supplemental Figure S4B-C). Interestingly in KD cells, we did not observe an accumulation of lipidated LC3 and GABARAP in the absence of BafA1 under starvation conditions (Figure 3G-H), BAG2 KD leading to an increase in autophagy flux (Figure 3I). This was also seen by employing the Halo-Tag reporter system (Yim et al., 2022) (supplemental Figure S4D-E), as well as by p62 puncta accumulation by IF (supplemental Figure S4F-G). Thus, whereas chronic removal of BAG2 leads to reduced autophagy flux, acute removal leads to an increase in autophagy flux. To address this difference, we monitored autophagosome-lysosome fusion by analyzing p62-LAMP2 colocalization by IF. Whereas we observed an increase in p62-LAMP2 colocalization in WT cells when lysosomal acidification was blocked by BafA1, and when autophagy was induced by starvation, we failed to observe this in BAG2 KO cells, indicating that lysosomal targeting of autophagosomal cargo is perturbed by the chronic loss of BAG2 (Figure 3J, supplemental Figure S5A). In addition, *BAG2* KO cells exhibited a perturbed lysosome localization under basal conditions, with a loss of a perinuclear lysosomal network. In shRNA-mediated BAG2 KD cells, p62-LAMP2 colocalization behaved similarly to control cells using a scrambled shRNA indicating that lysosomal targeting of autophagosomal cargo is not perturbed by the acute loss of BAG2 (Figure 3K, supplemental Figure S5B). Taken together, the acute effect of loss of BAG2 is an increase in autophagy flux. Under prolonged absence in KO cells, loss of BAG2 appears to have secondary effects that interfere with vesicle trafficking leading to a reduction in autophagosome-lysosome fusion and lysosomal miss-positioning, which leads to a net reduction of autophagy activity.

### BAG2 changes its intracellular localization/interaction partners in response to autophagy induction

Having established that BAG2 modulates autophagy, we next aimed to elaborate potential underlying mechanisms. For this, we expressed N-terminally HA-Streptactin (SHA)-tagged BAG2 in BAG2 KO cells by lentiviral infection and analyzed BAG2 interaction partners by AP-MS both in growth and in starvation conditions. In three biological replicates each, we identified 736 proteins as significantly enriched comparing SHA-BAG2-APs to respective negative controls i.e., doxycycline treated BAG2 knockout cells infected with a lentiviral control vector (total of 12 APs, q<0.05, supplemental Table S3A-C). This is a rather large number of potential interactors and likely reflects BAG2’s promiscuous function as HSC/HSP70 co-chaperone being involved in numerous cell biological processes. Of these enriched proteins, 97 bound stronger in starvation and 27 stronger in growth conditions (Figure 4A, q<0.05, supplemental Table S3C). Analyzing enriched GO terms to identify stimulus-dependent alterations, it appears that BAG2 changes its subcellular localization upon starvation. Whereas we identified several tubulin subunits as enriched under growth conditions, indicating a microtubular localization (Figure 4A), in autophagy-inducing conditions, enriched proteins carried GO terms related to endomembranes, specifically to vesicle coat and scaffold proteins, such as the COPII coat protein SEC13 and the ER exit sites (ERES) marker SEC16A (Figure 4A-B). Indeed, focusing on starvation-specific proteins which carry the GO term “endomembrane system” we were able to generate a protein interaction network of 46 proteins using STRING DB (Szklarczyk et al., 2023) (Figure 4C). In agreement with a change in subcellular localization to endomembranes upon starvation, a crude cell fractionation separating micro-vesicles, i.e. the 17’000 *g* fraction containing ER, Golgi, endo- and lysosomes (Dengjel et al., 2012), from cytosol, nuclei and mitochondria yielded an increased abundance of BAG2 in the RAB7-positive cell fraction in starvation (Figure 4D-E).

**Figure 4:**
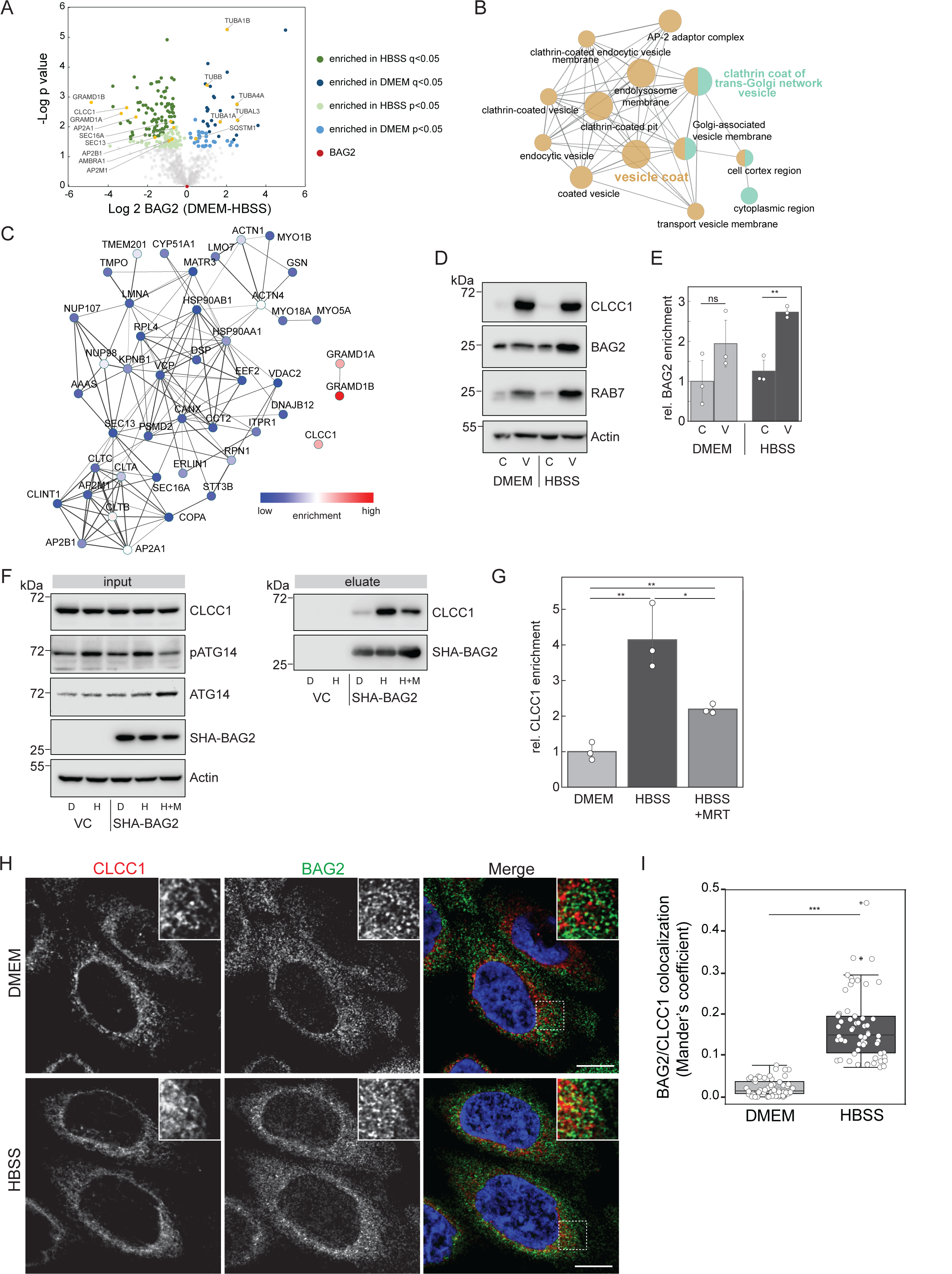
Upon starvation BAG2 changes its subcellular localization. **(A)** BAG2 interactome under growth (DMEM) and starvation conditions (HBSS). Hela cells inducibly expressing SHA-tagged BAG2 in comparison to vector ctrl. were used to determine stimulus-specific interaction partners by AP-MS. Proteins being significantly enriched compared to negative control experiments were analyzed on differential binding (n=3 biol. replicates each; Student’s t-test, unpaired, p<0.05, see figure for details). **(B)** BAG2 binds to proteins of endomembranes upon starvation. GO enrichment analysis of significant BAG2 interactors upon starvation. **(C)** Protein interaction network of starvation-specific BAG2 interactors. Color scale indicates level of enrichment. **(D-E)** Increased BAG2 abundance in microvesicular cell fraction under starvation. Cells were starved for 2 h (HBSS) or kept in growth conditions (DMEM), lysed by douncing, and fractionated into cytosol (C) and 17’000 *g* microvesicular fraction (V). In n=3 biological replicates a significant increase of BAG2 in the microvesicular fraction under HBSS conditions is observed. **: p<0.01, t-test. Single values are highlighted as white dots. Error bars indicate SD. **(F-G)** Upon starvation (2 h HBSS), BAG2 binds stronger to the chloride ion channel CLCC1. Treatment with 20 μM of the ULK1 inhibitor MRT68921 (2 h prior HBSS treatment, total of 4 h) partially blocks this increased interaction. (F) shows a representative biological replicate of n=3. Samples were normalized to protein amount prior analysis. (G) quantification of replicates shown in (F) normalized to BAG2 levels. *: p<0.05, **: p<0.01, t-test. Single values are highlighted as white dots. Error bars indicate 95% confidence interval. **(H-I)** Starvation increases the pool of ER-localized BAG2. (H) BAG2 colocalization with CLCC1 increases upon starvation. IF micrographs from untreated (DMEM) or starved (HBSS, 3 h) HeLa WT cells. Representative pictures of n=3 biological replicates are shown. Scale bar = 10 μm. (I) colocalization quantification of (H). Thresholded Mander’s coefficient B was calculated per cell and 60 cells of n=3 biological replicates. An unpaired student’s t-test was used to determine the significance of colocalization changes. ***: p<0.001. Error bars= SEM.

One of the most strongly binding proteins in starvation conditions was the ER/Golgi-localized chloride ion channel CLCC1, mutations of which have been associated with retinitis pigmentosa (Li et al., 2018) (Figure 4A, C). CLCC1 constitutively localized to the 17’000 *g* fraction, independent of growth conditions (Figure 4D). To further test if growth conditions lead to changes in intracellular localization and protein interactions of BAG2 and if this is dependent on ULK1 activity, we studied the BAG2-CLCC1 interaction in growth and starvation conditions in the presence and absence of the ULK1 inhibitor MRT68921 by AP coupled to western blot (Petherick et al., 2015). In starvation conditions, BAG2 interacts stronger with CLCC1 (Figure 4F-G), which is in agreement with MS analyses (Figure 4A); and this change in interaction is indeed dependent on ULK1 activity as MRT68921 treatment partially blocks it. Finally, also by IF, we observed that BAG2 colocalizes significantly stronger with both CLCC1 and SEC16A in starvation conditions (Figure 4H-I, supplemental Figure S6). SEC16A being a marker of ERES, one principal site of autophagosome biogenesis (Graef et al., 2013; Tan et al., 2013). To further characterize the CLCC1-positive membrane compartment, its co-localization with membrane-bound organelle markers was analyzed by IF. We observed partial co-localization of CLCC1 with LMAN1/ERGIC-53, SEC13, but not with SNX1 indicating that CLCC1 at least partially localizes to the ER-Golgi intermediate compartment (supplemental Figure S7). Taken together, under starvation i.e., autophagy induction, a pool of BAG2 changes its subcellular localization and protein-protein interactions in an ULK1-dependent manner localizing to CLCC1-positive endomembranes, which include membranes that are positive for SEC13 and SEC16A being indicative for ERES, sites at which autophagosome biogenesis takes place.

### In nutrient starvation BAG2 supports AMBRA1 recruitment to the ER

One of the proteins that we found reproducibly significantly enriched in starvation conditions by MS-based proteomics of exogenous and endogenous BAG2 APs and that has a known function in autophagy regulation is the VPS34 complex member AMBRA1 (Antonioli et al., 2017; Cianfanelli et al., 2015) (Figure 4A, 5A). In agreement, we also identified an increased interaction between AMBRA1 and BAG2 in starvation conditions by comparing APs of tagged BAG2 using western blot (Figure 5B-C) as well as by IPs of endogenous AMBRA1 using MS as readout (Figure 5D). Thus, AMBRA1 binds to BAG2 and this binding is increased by limited nutrient availability. The interaction of AMBRA1 with BAG2 might be direct or indirect via HSC/HSP70 members. To discriminate these two options, we expressed the HSC70-binding deficient BAG2 variant BAG2^I160A^ (Carrettiero et al., 2022) in BAG2 KO cells and compared its interactome to BAG2^WT^ expressing cells by AP-MS. As anticipated, HSPA4 and HSPA9 bound weaker to BAG2^I160A^ (Figure 5E), as did CLCC1 indicating that the latter might interact indirectly with BAG2 via HSC70 proteins. Interestingly, AMBRA1 bound equally well to both BAG2 variants implying that this interaction is potentially direct and independent of HSC/HSP70-BAG2 interactions (Figure 5E).

**Figure 5:**
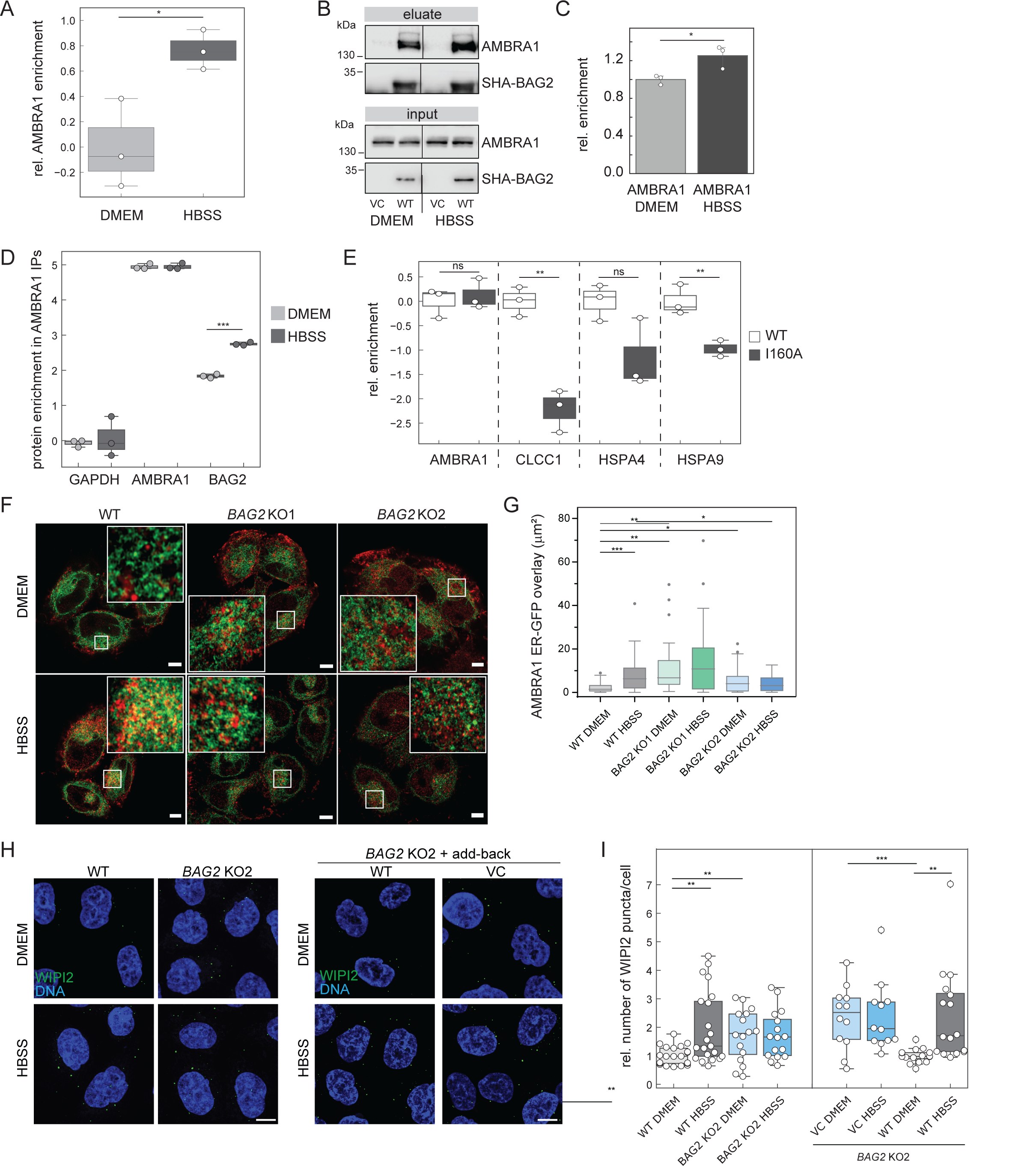
BAG2 regulates the subcellular distribution of AMBRA1. **(A)** AMBRA1 binds stronger to SHA-BAG2 under starvation conditions. Shown is the relative enrichment of AMBRA1 in BAG2-AP as determined by MS of n=3 biological replicates. Box plots show spreads of log2 transformed MS ratios compared to neg. control experiments. Single values are highlighted as white dots. Error bars indicate 95% confidence intervals. **(B-C)** AMBRA1 binds stronger to SHA-BAG2 under starvation conditions as determined by AP-western blot analysis. (B) shows one representative western blot of n=3 biological replicates. Samples were normalized to protein amount prior analysis. Note: samples were run on one gel, black lines indicate cropping of unrelated lanes. (C) quantification of replicates shown in (B). *: p<0.05, t-test. Error bars indicate 95% confidence interval. **(D)** Endogenous AMBRA1 and BAG2 interact. Shown are relative abundances of annotated proteins in AMBRA1 IP-MS experiments. Starvation (1 h HBSS) leads to an increase in AMBRA1-BAG2 interaction as compared to growth conditions (DMEM). GAPDH is shown as negative control. Box plot shows log2-transformed relative fold changes of protein abundances (iBAQ values) compared to IgG control experiments normalized to AMBRA1 enrichment. Error bars indicate 95% confidence interval. Single values of n=3 biological replicates are highlighted as white dots. ***: p<0.001. **(E)** The interaction of AMBRA1 with BAG2 does not depend on HSC/HSP70 binding. Shown are relative abundances of annotated proteins in SHA-BAG2 AP-MS experiments. Compared to BAG2^WT^ (WT), BAG2^I160A^ (I160A) interacts less with CLCC1, HSPA4 and HSPA9. The interaction with AMBRA1 is not perturbed by the point mutation. Box plots show log2-transformed relative fold changes of protein abundances (iBAQ values) normalized to WT, error bars indicate 95% confidence interval. Single values of n=3 biological replicates are highlighted as white dots. **: p<0.01; ns: not significant. **(F-G)** Loss of BAG2 leads to perturbed recruitment of AMBRA1 to ER membrane. (F) Representative fluorescence images of ERGFP expressing HeLa WT and BAG2^KO/KO^ cells stained for AMBRA1 (Alexa 647) and subjected, or not, to nutrient deprivation, adding HBSS for 2 h. Scale bar: 5 μm. (G) Quantitative analysis of GFP and Alexa 467 overlayed area (μm^2^) per cell reported as the mean ± SD, *P ≤ 0.05, **P ≤ 0.005 and ***P ≤ 0.0005 using ANOVA 2-way test for repeated samples. **(H-I)** Loss of BAG2 leads to increased number of WIPI2 dots. (H) WIPI2 puncta analysis using IF micrographs from untreated (DMEM) or starved (HBSS, 3 h) HeLa WT and BAG2 KO cells (left side), or BAG2 KO cells transfected with a lentiviral control vector (VC) or a vector encoding BAG2^WT^ (WT, right side). Representative pictures of n=3 biological replicates are shown. Scale bar = 10 μm. (I) Puncta quantification of (H). White dots represent the quantified average WIPI2 puncta per cell per image (5 to 23 cells per image, with a total of 12 images of n=3 biological replicates for BAG2 KO + VC samples, or 16 images of n=4 biological replicates for each other conditions). An unpaired student’s t-test was used to determine the significant differences. **: p<0.01; ***: p<0.001.

To address the relevance of the BAG2-AMBRA1 interaction, we studied the subcellular localization of AMBRA1 in HeLa WT and BAG2 KO clones by IF. In the absence of BAG2 we observed a perturbed intracellular distribution of AMBRA1. Whereas one could observe an increased localization of AMBRA1 to the ER membrane in starvation in wildtype cells, KO cells lost their ability to respond to starvation and exhibited an increased localization of AMBRA1 to the ER membrane also in growth conditions (Figure 5F-G). This implies that one role of BAG2 in autophagy is the regulation of the subcellular distribution of AMBRA1, sequestering AMBRA1 in growth conditions at microtubuli (Di Bartolomeo et al., 2010), and interfering with its localization to the ER membrane. As both KO clones behaved similarly, we performed further experiments with clone 2 only. To analyze if the observed changes in AMBRA1 localization mirror changes in VPS34 complex activity affecting the local production of PI3P, we monitored WIPI2 puncta formation in WT and BAG2 KO cells. WIPI2, an ortholog of yeast Atg18 is a PI3P-binding protein important for autophagosome expansion and serves as a docking site for the ATG5-ATG12/ATG16 complex (Dooley et al., 2014). Its local accumulation in puncta can be used as proxy to address VPS34 activity (Axe et al., 2008; Polson et al., 2010). Reflecting the observation in ER-AMBRA1 localization, we also identified an increase in WIPI2 puncta in BAG2 KO cells in growth conditions that did not further increase under starvation conditions, in contrast to WT cells (Figure 5H-I). This increase was dependent on the absence of BAG2 as its ectopic re-expression in KO cells rescued this phenotype in contrast to KO cells expressing a control vector (Figure 5H-I). We observed the same phenotype in cells acutely depleted of BAG2 by the expression of a doxycyclin-inducible shRNA (supplemental Figure S8). Thus, our results imply that acute or chronic loss of BAG2 leads to an increase in WIPI2-positive autophagosome nucleation sites, which is mirrored by an increased lipidation of LC3 and GABARAP under basal growth conditions in KO cells (Figure 3D-E).

### BAG2 Ser31 is a ULK1 target site regulating its protein interactions and subcellular localization

Finally, we addressed the question if ULK1 may directly regulate localization and/or protein-protein interactions of BAG2 by phosphorylation. In a phosphoproteomic screen, we identified Ser31 of BAG2 as a potential ULK1 target site (Hu et al., 2021). Ser31 lies within a predicted coiled-coil region, is highly conserved in mammals, and the surrounding amino acids fit to the ULK1 consensus motif with a leucine in the −3 position (Figure 6A) (Egan et al., 2015). To test whether ULK1 indeed directly phosphorylates BAG2 at amino acid residue Ser31, we performed filter-aided *in vitro* kinase assays coupled to MS-based quantification of phosphopeptides (Hu et al., 2019). Briefly, BAG2 purified from HeLa cells was dephosphorylated by lambda phosphatase and incubated with purified ULK1^WT^ or a kinase dead-variant of ULK1 (ULK1^KD^), in which Asp165 was replaced by Ala (Loffler et al., 2011), phosphopeptides were enriched and quantified by LC-MS/MS using extracted ion currents. Whereas we identified the tryptic peptide carrying the Ser31 phosphorylation site in two biological replicates using ULK1^WT^ as kinase, we failed to identify this site in *in vitro* kinase assays using ULK1^KD^ (Figure 6B). Non-phosphorylated peptides of BAG2 were present in similar amounts. Thus, Ser31 of BAG2 fulfils all requirements of being a *bona fide* ULK1 target site.

**Figure 6:**
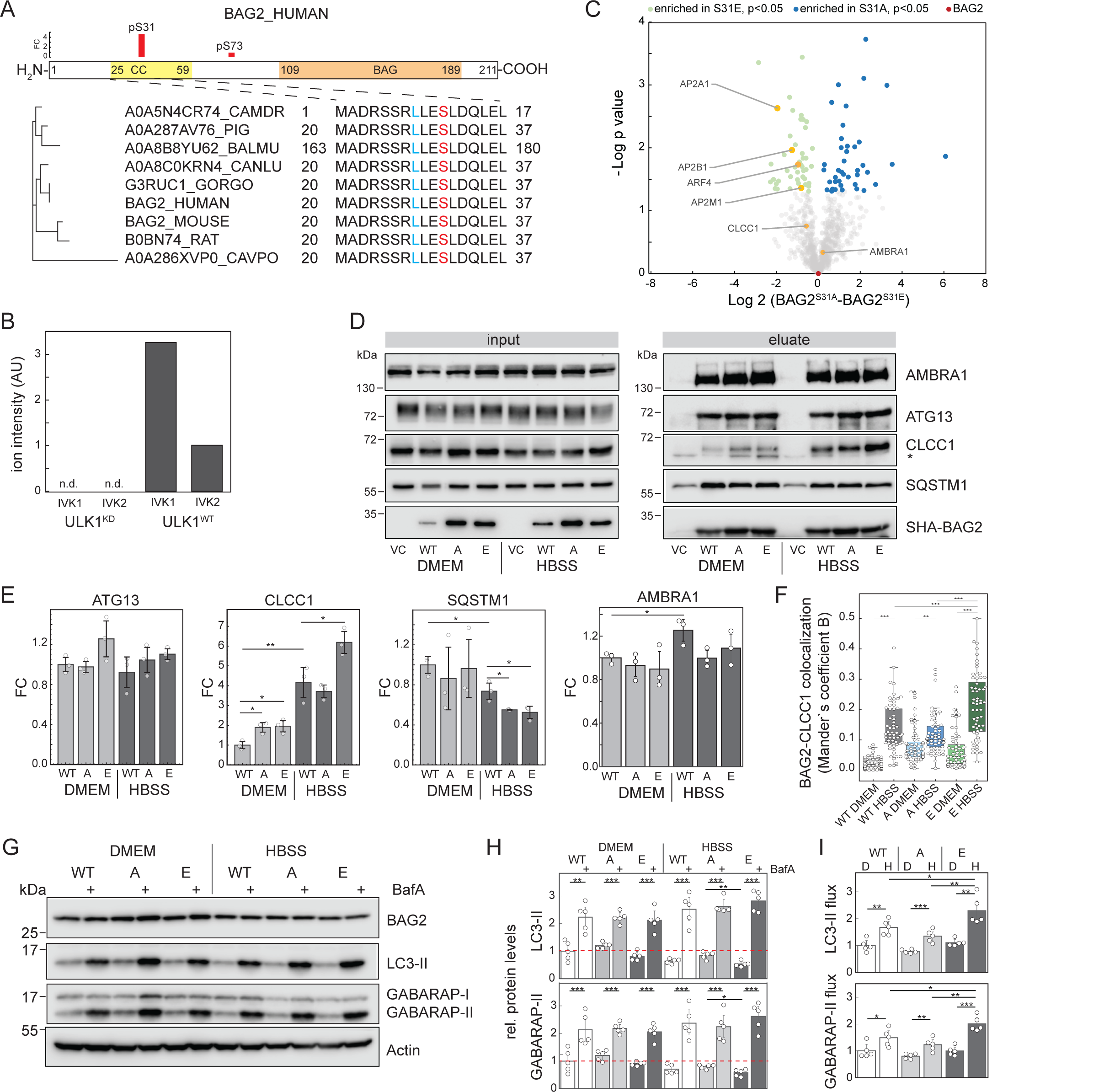
BAG2 is a ULK1 target, its phosphorylation affecting localization and protein-protein interactions. **(A)** BAG2 Ser31 is an *in vivo* ULK1 target site. Ser31 was identified as being significantly phosphorylated in MEFs expressing human ULK1 compared to ULK1 KO MEFs, in contrast to Ser73 which did not differ depending on the ULK1 status (Hu et al., 2021). Red bars indicate fold change of phosphorylation (n ≥ 2 biol. replicates). Ser31 lies within a ULK1 consensus motif which is highly conserved in mammals. (CAMDR: Camelus dromedarius, Arabian camel; PIG: Sus scrofa, Pig; BALMU: Balaenoptera musculus, Blue whale; CANLU: Canis lupus, Gray wolf; GORGO: Gorilla gorilla gorilla, Western lowland gorilla; CAVPO: Cavia porcellus, Guinea pig). **(B)** BAG2 Ser31 is phosphorylated by wildtype ULK1 (ULK1^WT^) *in vitro,* but not by kinase-dead ULK1 (ULK1^KD^). Shown are two biological replicates (IVK1, IVK2). **(C-E)** ULK1 Ser31 status affects BAG2-protein interactions. (C) volcano plot of AP-MS data highlighting differential binding of BAG2 Ser31 variants. (D) Co-IP western blot of SHA-tagged WT-, Ser31Ala (A)- and Ser31Glu (E)- BAG2 variants. Shown is one representative of n=3 biological replicates. (E) Quantification of replicates shown in (D). Samples were normalized to protein amount prior analysis. Error bars indicate 95% confidence interval. Single values are highlighted as white dots. *: p<0.05, **: p<0.01, unpaired T test. Note: Blots shown in panel D were also used in Figure 3B; quantifications of WT bands were also shown in Figures 3C, and 5C. **(F)** Expression of BAG2 Ser31Glu (E) leads to a significant increase in colocalization of BAG2 and CLCC1 compared to BAG2 WT and BAG2 Ser31Ala (A) expressing cells in starvation conditions (3 h, HBSS) as analyzed by IF. Data of n=3 biological replicates each, 20 cells per replicate. White dots represent the colocalization foci count per cell and per biological replicate. An unpaired two-tailed Student’s t-test was used to compare values. *: p **≤** 0.05, **: p **≤** 0.01, ***: p **≤** 0.001. Respective immunofluorescence micrographs are shown in supplemental Figure S9. **(G-I)** Cells expressing BAG2 Ser31Glu (E) exhibit an increased autophagy flux in starvation compared to BAG2 WT and BAG2 Ser31Ala (A) expressing cells. (G) Shown are representative western blots of n=5 biological replicates. Actin serves as loading control. (H) Quantifications of blots shown in (G). Red dotted lines indicate protein levels in DMEM control conditions. (I) Comparing BafA1-treated with non-treated lanes indicates increased autophagy flux in starvation (HBSS, H) compared to growth conditions (DMEM, D). BAG2 Ser31Glu (E) leads to a significant increase in autophagy flux in starvation compared to BAG2 WT and BAG2 Ser31Ala (A). Student’s t-test, unpaired, *: p<0.05, **: p<0.01, ***: p<0.001. Single values are highlighted as white dots. Error bars indicate 95% confidence interval.

To address the functional relevance of the identified phosphosite, we generated a non-phosphorylatable Ala variant (BAG2^S31A^) and a phosphomimicking Glu variant (BAG2^S31E^) of BAG2 and expressed them in BAG2 KO cells. Both, by AP-MS as well as by AP-western blot we observed increased interaction of CLCC1 with BAG2^S31E^ under starvation conditions (Figure 6C-E, supplemental Table S4). In contrast, the interactions of BAG2 with AMBRA1 and ATG13 were not affected by the site variants, whereas the interaction with p62was significantly decreased (Figure 6D, E). In agreement with the presented biochemical data, also IF analyses of cells expressing the various BAG2 variants revealed a significantly increased colocalization between BAG2^S31E^ and CLCC1 under starvation conditions in contrast to BAG2^S31A^ (Figure 6F, supplemental Figure S9). These data imply that ULK1 phosphorylation of BAG2 at Ser31 regulates its interaction with the transmembrane protein CLCC1 which reflects a ULK1-dependent change in subcellular localization.

Finally, we also analyzed if the BAG2 variants differentially affect the observed autophagy phenotype in BAG2 KO cells. In starvation conditions, expression of *BAG2^S31E^* but not *BAG2^S31A^*led to increased autophagy flux compared to *BAG2^WT^*as analyzed by LC3-II and GABARAP-II lipidation (Figure 6 G-I). Thus, ULK1 phosphorylation of BAG2 at Ser31 supports localization of BAG2 to CLCC1-positive endomembranes conveying a positive effect to autophagy activity.

## Discussion

Autophagy is a vital catabolic pathway; its dysregulation being implicated in numerous human diseases (Klionsky et al., 2021). The ULK1 kinase complex is crucial for canonical autophagy regulation and represents an interesting drug target which might support the specific modulation of autophagy in diverse disease settings (Whitmarsh-Everiss and Laraia, 2021). In the current manuscript we present a deep ULK1 interactome using a combination of AP- and PL-MS-based proteomics. By employing N- and C-terminally tagged protein variants we characterize 317 high confidence subunit-specific and common ULK1-complex interactors. Focusing on BAG2, a common interactor that bound to more than one subunit and was identified by AP- and PL-MS, we characterized it as a new ULK1 effector protein that modulates functional autophagy by regulating the subcellular localization of AMBRA1.

MS-based proteomics is indispensable for the unbiased identification of protein-protein interactions and PTMs. The introduction of PL approaches supports the characterization of *in vivo* protein neighborhoods not necessarily focusing on direct interactors but rather on proteins being enriched in the vicinity of bait proteins. Compared to classical AP-MS approaches neighboring proteins are marked by chemical tags in their native environment, which reduces post-lysis artefacts (Branon et al., 2018). Thus, AP- and PL-MS are orthogonal approaches that commonly lead to the identifications of different sets of target proteins (Lambert et al., 2015; Liu et al., 2020). Also in the current study, the overlap between interactors identified by AP- and PL-MS was rather small. What came as surprise was the observation that PL-MS yielded a smaller number of neighbors compared to AP-MS-based interactors when the bait protein has a similar size as the tag itself. Assuming that this is a general trend and not specific to ATG101, a positive interpretation of this observation is that smaller proteins might be more mobile, PL leading to more noise and by this interfering with the identification of transiently interacting proteins. A negative interpretation would be that the relative size of the tag has a substantial influence on protein localization, large tags leading to miss-localized i.e., non-physiological localized proteins. As we used a double-tag strategy combining both the PL-enzyme miniTurbo with the HA-peptide epitope in single fusion proteins, we tend to support the first interpretation. In our case, protein miss-localization should also be reflected by altered/reduced stable interactions identified by AP-MS, which was not the case.

The presented deep ULK1 complex interactome is a rich resource for research groups being interested in autophagy and its regulation. We highlighted the interaction with other protein and lipid kinases and phosphatases leading to the interpretation that the ULK1 complex forms a signalosome. This is further supported by the observation that autophagy initiation leads to clustering of ULK1 complex members at ER-sub-compartments which is observable by fluorescence microscopy i.e., ULK1 or ATG13 puncta formation at omegasomes (Karanasios et al., 2013; Karanasios et al., 2016). In agreement, it was recently shown that under starvation conditions ULK1 forms large clusters at the ER containing more than 100 molecules (Banerjee et al., 2023). Interestingly, we identified several SARs as common ULK1 complex interactors under starvation conditions. Nutrient starvation, in our case the lack of amino acids, growth factors and lipids (the only remaining energy source being glucose), is the classical stimulus for non-selective bulk autophagy. The enrichment of SARs implies that selective autophagy sub-pathways are also active under “bulk” starvation conditions. Indeed, non-selective bulk autophagy could also be envisioned as the sum of many selective autophagy pathways happening in concert (Kirkin and Rogov, 2019; Nair and Klionsky, 2005; Zaffagnini and Martens, 2016). Time-resolved, detailed characterizations of molecular events and autophagosomal proteome compositions are needed to further disentangle selective from potential bulk pathways (Kallergi et al., 2023).

Having performed “omics” type of experiments always raises the questions which and how many of the identified targets should be studied and to which detail. We decided to follow-up a single ULK1 interactor, the HSP70 co-chaperone BAG2, and to analyze the events that govern its function in autophagy in molecular detail. In contrasts to other members of the BAG protein family (Pattingre and Turtoi, 2022), the primary function of BAG2 appears to be the inhibition of autophagy under basal growth conditions, as its acute depletion by shRNA-based KD leads to increased autophagy flux. The increased flux is due to increased autophagosome nucleation sites as indicated by WIPI2 puncta formation. The increased number of WIPI2 puncta under basal conditions also indicates that the inhibitory function of BAG2 is likely independent of ULK1 activity. In KO cells, the phenotype is more complex: in agreement with the observations in KD cells, increased lipidation of ATG8 proteins and increased WIPI2 puncta numbers are observed; however, in contrast to the effect of acute reduction of BAG2 abundance by shRNA, the chronic reduction of BAG2 levels by KO leads to reduced autophagy flux as analyzed by western blot, MS and IF. That latter appears to be an indirect effect due to perturbation of lysosomal targeting related to loss of BAG2 as shown by reduced colocalization of p62 and LAMP2. Indeed, we identified BAG2 interacting with several vesicle coat proteins: clathrin-coat-proteins like adapter protein complex 2 members AP2A1, B1 and M1 which are important for clathrin-coated vesicle formation and cargo selectivity (Owen et al., 2004), and to COPII-coat proteins like SEC13. Hence, BAG2 might have a general effect on vesicle formation and/or trafficking affecting autophagosome turnover also in an indirect manner. Further studies are needed to fully uncover the role of BAG2 in intracellular vesicle trafficking, fusion, and lysosomal activity. Under starvation conditions and activation of ULK1, BAG2 changes its protein interactions and with this its subcellular localization in a ULK1-dependent manner, localizing more to endomembranes, as exemplified by binding stronger to the ER-/Golgi-resident transmembrane protein CLCC1. In a recent spatial proteomics approach, BAG2 was identified as localizing to SEC16A-positive ER exit sites (Nalbach et al., 2023), which are involved in autophagosome biogenesis (Yang et al., 2021). Our data recapitulate this finding additionally implying that BAG2 may localize to different membrane subcompartments, which would support the interpretation that BAG2 may fulfill multiple functions important for autophagy activity. As the *C. elegans* homologs of BAG2 and HSC70 have been shown to regulate intracellular protein localization (Fukuzono et al., 2016), we hypothesize that the change in BAG2 localization likely affects also BAG2 client proteins. One autophagy-relevant client of BAG2 that was identified in the current study is AMBRA1.

KO of BAG2 supported increased AMBRA1 localization to the ER membrane. The increase in WIPI2 puncta in both KO and KD cells is in agreement with ER-localized AMBRA1, as their formation is PI3P dependent reflecting an increase in VPS34 kinase activity (Axe et al., 2008; Polson et al., 2010). If the increase in autophagosome initiation sites in BAG2 depleted cells represent ER-connected phagophores or accumulated ATG9 vesicles will have to be addressed in future studies (Broadbent et al., 2023; Cook and Hurley, 2023; Olivas et al., 2023). The change in BAG2 localization is directly regulated by ULK1, as pharmacological inhibition of ULK1 decreased and mutating the ULK1 target site Ser31 of BAG2 to a phosphomimetic glutamic acid increased CLCC1 binding, respectively. The respective alanine mutation had no effect. Indeed, by using *in vivo* and *in vitro* phosphoproteomics, we show that Ser31 of BAG2 is a *bona fide* ULK1 target site. Interestingly, the effects of *BAG2^S31E^* expression and starvation seem to be additive with respect to CLCC1 binding implying that the interaction does not solely rely on Ser31 phosphorylation. Additional amino acids of BAG2 or CLCC1 might be modified in the tested condition affecting binding, either via phosphorylation by ULK1 or other kinases, or via additional PTMs, potentially also affecting AMBRA1 binding. The fact that the BAG2-CLCC1 interaction is also dependent on HSC70 binding as exemplified by the effects of the BAG2 Ile160Ala mutation further supports the interpretation that next to Ser31 phosphorylation additional mechanisms affect this interaction. Further studies are needed to characterize the regulation of BAG2 and the immediate effect of BAG2 Ser31 phosphorylation. It is also not clear how the interaction between AMBRA1 and BAG2 is regulated on a molecular level.

The synonym of AMBRA1 is DCAF3 (DDB1- and CUL4-associated factor 3) as AMBRA1 has also been identified as a binding partner of E3 ubiquitin ligases, amongst them Cullin 4 and Cullin 5 (Antonioli et al., 2014; Jin et al., 2006), TRAF6 (Nazio et al., 2013), and TRIM32 (Di Rienzo et al., 2019). Indeed, AMBRA1 and ULK1 appear to be linked by feedbacks, ULK1 phosphorylating AMBRA1 (Di Bartolomeo et al., 2010) which in turns supports the TRAF6-dependent ubiquitination and stabilization of ULK1 (Nazio et al., 2013). Whereas we did not observe clear BAG2 abundance changes comparing growth and starvation conditions, we cannot exclude that AMBRA1 affects the ubiquitination of BAG2 and with this its stability or protein-protein interactions, affecting potentially both autophagosomal and proteasomal protein degradation. This will have to be addressed in the future.

Taken together, in the current study we generated a deep ULK1 complex interactome highlighting that the holo-complex forms an elaborate signalosome. By phosphorylating the co-chaperone BAG2, ULK1 modulates its subcellular localization, positively affecting autophagy initiation in response to stress by stabilizing AMBRA1 at the ER membrane. However, next to regulation of autophagy initiation, the ULK1 holo-complex appears to modulate adaptation to limited nutrient supply in general. SARs are important binding partners of ULK1 complex members indicating the activation of starvation-dependent selective autophagy pathways and questioning the relevance of non-specific bulk degradation by autophagy in response to stress.

## Materials and Methods

### Cell Culture and treatments

HeLa cells (ATCC CCl-2) were validated by genotyping (Microsynth) and negatively tested for mycoplasma. All cells were grown in a humidified incubator in Dulbecco’s modified Eagle medium (DMEM, PAN Biotech, P04-04510) supplemented with 10% fetal bovine serum (FBS, BioWest, S181B-500), 1% penicillin/streptomycin (PAN Biotech, P06-07100) at 37°C and 5% CO2. For SILAC labeling, cells were cultured for seven generations using SILAC-DMEM, containing non-labeled and labeled lysine and arginine variants: Lys-8 (Sigma, 608041) and Arg-10 (Sigma, 608033) for heavy, Lys-4 (Sigma, 616192) and Arg-6 (Sigma, 643440) for medium, and Lys-0 (Sigma, L8662) and Arg-0 (Sigma, 11039) for light labels, supplemented with 10% dialyzed FBS (Biowest, S181D-500)1% penicillin/streptomycin and 1% glutamax (GIBCO, 35050038). The final concentration of arginine was 42 mg/L, lysine of 73 mg/L and to avoid the conversion of arginine to proline, L-proline 26 mg/L was added. For phenotype assays, HA affinity purification, and cellular fractionation experiments, starvation was induced by incubating cells with Hank’s Balanced Salt solution (HBSS, ThermoFisher, 14025-100) for 2 h. For proximity labeling experiments and HA affinity purification of ULK1 complex members, cells were incubated with HBSS for 90 mins. In all indicated experiments 10 nM concanamycin A (Sigma-Aldrich, C9705) or 2 nM Bafilomycin A1 (Santa Cruz Biotechnologies, sc-201550A) were used. ERGFP stably expressing HeLa cells were obtained as previously described (Di Bartolomeo et al., 2010).

### Cloning of recombinant DNA constructs

The following gene-encoding plasmids were generous gifts from Prof Björn Stork: ULK1 WT, ULK1 KD (kinase dead, D165A), ATG13 isoform II, and FIP200/RB1CC1. BAG2 and ATG101 were amplified by RT-PCR from HEK293T cDNA (Qiagen QuantiTect RT-PCR kit, 205311) using primers oNDS227 and oNDS228 for ATG101, and oNDS309 and oNDS310 for BAG2 (please see table: S2A for all primer sequences). The GFP gene was PCR amplified using primers oNDS139 and oNDS140 from pEGFP-C1-puromycin plasmid to make the miniTurbo-GFP fusion (pDS48) plasmid. miniTurbo containing plasmid 3xHA-miniTurbo-NLS_pCDNA3 was a gift from Alice Ting (Addgene plasmid # 107172). miniTurbo gene (primers: oNDS:115 and 116) was fused N-terminal to all constructs (ULK1 (pDS38) (oNDS:117 and 118), FIP200 (pDS41) (oNDS:121 and 122), ATG13 (pDS40) (oNDS:119 and 120) and ATG101 (pDS42) (oNDS:231 and 232)) starting with N-terminal HA-tag, miniTurbo, V5-tag and the gene of interest. Similarly, for the C-terminal tagging, the sequence starts with N-terminal gene constructs (ULK1 (pDS43) (oNDS:190 and 191), FIP200 (pDS46) (oNDS:188 and 189), ATG13 (pDS45) (oNDS:225 and 226) and ATG101 (pDS47) (oNDS:229 and 230)), followed by V5-tag, miniTurbo (oNDS:132 and 133) and HA tag at the C-terminal end. A PCR fragment of miniTurbo containing HA- and V5-tag was generated and another PCR product for gene of interest (please see table: S1A for primer sequences) having the V5-tag and homology arms with the vector was generated. All PCR reactions were performed using the Phusion high fidelity DNA polymerase (NEB, M0530S). All the PCR-amplified fusion genes and epitope tags were cloned into the pSKP32 vector (Kaeser-Pebernard et al., 2022) between the NheI and PstI restriction sites using Gibson assembly (NEB, M5510A) according to manufacturer’s protocol. Using the same method, HA-2X-Streptactin tag (SHA, (primers: oNDS107/111)) was fused to the N-terminal of BAG2 WT (pDS49) (primers: oNDS318/319), phospho-mutants (BAG2-S31A (pDS50) (primers: oNDS345/348/346/347), and S31E (pDS51) (primers: oNDS345/348/349/350) genes and cloned into pSKP32. HSC/HSP70 deficient mutant BAG2 I160A (pDS52) (primers: oNDS345/348/379/380) was also cloned the same way in pSKP32. The pSKP32 vector was used as a negative/beads control for all SHA experiments. A pSKP190 destination vector was generated by replacing the hPGK-Puromycin resistance cassette of pCW57.1 (gift from David Root (Addgene plasmid # 41393; http://n2t.net/addgene:41393; RRID:Addgene_41393)) between the AgeI/XbaI restriction sites with a mPGK-Hygromycin resistance cassette PCR amplified from pER81 (generous gift from Richard Iggo) using oSKP-286/287 by Gibson assembly. The inducible Halo-rLC3B expression lentiviral vector pSKP199 was generated as follows. The Halo tag gene was purchased from DNASU (HsCD00840399), PCR amplified (primers: oSKP 298/299), andprecombinant rat LC3B (rLC3B) was synthesized as a double stranded gene block (IDTDNA). pSKP190 was double digested with NdeI/BsrGI. Gibson assembly was performed by assembling Halo tag, rLC3B (for sequence please see S2F) and the double digested pSKP190 vector. All the cloning constructs were sequence confirmed (Eurofins) with the listed primers (please see table: S2B for primer sequences).

### Cloning of CRISPR Cas-9 knockout vectors and cell line generation

CRISPR knockouts of BAG2, ULK1, FIP200, ATG13 and ATG101 in HeLa cells were performed using sgRNAs (single guide RNAs, please see table: S2C for guide RNA sequences) cloned in pX330-U6-Chimeric_BB-CBh-hSpCas9 (gift from Feng Zhang (Addgene plasmid # 42230; http://n2t.net/addgene:42230; RRID:Addgene_42230)). The sgRNAs were designed using CHOPCHOP CRISPR guide RNA algorithm (Labun et al., 2019), CRISPR plasmids and pEGFP-C1 puromycin plasmids were co-transfected in HeLa cells using lipofectamine LTX reagent (ThermoFisher, 15338030), according to manufacturer’s instructions. 24h post-transfection, co-transfected cells were selected in culture medium supplemented with 3 μg/ml puromycin (Invivogen, ant-pr-1), until selection was complete after about 48 h. Efficiently transfected cells were isolated to generate clonal lineages by single-cell cloning in 96-well plates; colonies were all evaluated for KO efficiency by western blot against the targeted protein. shRNAs were obtained from NEXUS Personalized Health Technologies (ETH Zürich). Scrambled shRNA as a control and shRNABAG2 (TRC0000033591, for sequence see table S2C) chosen from the Broad RNAi consortium were used for knockdown experiments. The above sequences were cloned between AgeI and EcoRI sites in pLKO.1 vector containing puromycin resistance gene leading to constitutive expressing. All cloning constructs were sequence confirmed using primer oNDS317.

### Lentivirus production and infection of target cells

To produce replication-defective lentiviral particles, recombinant DNA constructs cloned in lentiviral plasmids described above were co-transfected with packaging and envelope plasmids psPAX2 (gift from Didier Trono (Addgene plasmid # 12260; http://n2t.net/addgene:12260; RRID:Addgene_12260)) and pMD2.G (gift from Didier Trono (Addgene plasmid # 12259; http://n2t.net/addgene:12259; RRID:Addgene_12259)) into HEK293T cells seeded the night before using JetPrime transfection reagent (Polyplus, 114-75). Transfection medium was changed 12 h post transfection, and lentiviral supernatants were harvested 24 h later, sterile filtered through 0.2 µm syringe filters, supplemented with 8 µg/ml polybrene (Sigma-Aldrich, H9268), and stored in aliquots at −80°C. Lentiviral particles having different miniTurbo fusion constructs were used to infect respective HeLa ULK1 KO, ATG13 KO, FIP200 KO, and ATG101 KO cells. SHA-BAG2, BAG2 phospho-variants and corresponding vector control viral particles were used to infect the indicated BAG2 KO clones. Halo-rLC3B tag lentiviral particles were used to infect HeLa WT and BAG2 KO clones. To infect, viral dilutions (1:2 to 1:1000) were used in DMEM supplemented with 8 µg/ml polybrene. Infected cells were selected 24 h post infection in 4 µg/ml Blasticidin (Invivogen, ant-bl-1) or 100 µg/ml of hygromycin (Invivogen, ant-hg-1), according to the respective lentiviral selection marker, until selection was complete. Selected cells were tested for ectopic expression using doxycycline (2 μg/ml) inducible promoters by western blotting against the respective proteins-of-interest using anti-HA antibodies to determine the best working dilution. For knock-down experiments, shRNA-expressing lentiviruses were generated and used to infect the indicated target cells with the same protocol. 24 h post-infection, cells were selected in culture medium supplemented with 3 µg/ml puromycin (Invivogen, ant-pr-1), and selection was completed for 48 h before cells were harvested. Western blot analysis against BAG2 protein was performed using cell lysates to check efficiencies of shRNAs and the best shRNA was chosen for all shRNA related experiments. For shRNA experiments in Halo-rLC3B-expressing cells, a similar protocol was applied to the exception of Halo-rLC3B induction that was performed 24 h before TMR pulsing and harvesting.

### Antibodies

For western blotting experiments, all primary antibodies were used at a concentration of 1:1000 in either 5% milk or 5% BSA in 1X TBST buffer (1X TBS (Tris buffered saline), 0.1% Tween 20). Secondary antibodies were used at dilutions of 1:10’000 for anti-rabbit and 1:5’000 for anti-mouse in 5% milk or 5% BSA in 1X TBST. All primary and secondary antibodies were tested according to the manufacturer’s protocols. The following primary antibodies were used: Anti-β-Actin (Santa Cruz Biotechnologies, sc-47778, mouse monoclonal), SQSTM1/P62 (Cell Signaling, 5114S, rabbit), LC3A/B (Cell Signaling, 4108S, rabbit polyclonal), ATG14 (Cell Signaling, 96752, rabbit), ATG14-p29 (Cell signaling, 92340S, rabbit), FIP200 (Cell Signaling, 12436S, rabbit), ULK1 (Cell Signaling, 8054, rabbit), ULK1-p757 (Cell Signaling, 6888S, rabbit), ATG13 (Sigma, SAB4200100, rabbit), ATG101 (Cell Signaling, 13492, rabbit), BAG2 (Santa Cruz, sc-390107, mouse), BAG3 (Santa Cruz, sc-136467, mouse), BAG5 (Santa Cruz Biotechnologies, sc-390832, mouse), CLCC1 (Atlas, HPA013210, rabbit), WIPI2 (BioRad, MCA5780GA, mouse), AMBRA1 (Santa Cruz Biotechnologies, sc-398204, mouse), RAB7 (ABcam, ab137029, rabbit), Halo-Tag (Promega, G9211, mouse), and HSP90 (Santa Cruz Biotechnologies, sc-13119, mouse). Secondary antibodies used in the western blots were: Peroxidase conjugated anti-mouse monoclonal IgG (Jackson Immuno research, 111-035-062, RRID: AB_2338504), Peroxidase conjugated anti-rabbit monoclonal IgG (Jackson Immuno research, 111-035-045, RRID: AB_2337938)

For immunofluorescence (IF) experiments, all primary antibodies were used at a 1:100 dilution. In addition to those mentioned above, the following primary antibodies were used only in IF experiments: BAG2 (Novus Biologicals, NBP2-59476, mouse), ERGIC53 (Enzo, ENZ-ABS300-0100, mouse), SNX1 (BD Biosciences, 611482, mouse). Secondary antibodies used for IF were: donkey anti-mouse IgG (H+L) Alexa Fluor 488 (ThermoFisher, A21202), goat anti-rabbit Alexa Fluor 633 (ThermoFisher, A21071).

### Immunostainings and florescence microscopy analysis

For immunofluorescence analysis, the indicated cells were seeded on collagen I (ThermoFisher, A10483-01, diluted in 0.02 M acetic acid to 50 μg/ml)-coated coverslips for 24 h prior to experiments. Cells were treated as indicated and further fixed using either ice-cold 100% methanol (MetOH) or PFA (paraformaldehyde) for 15 min. Before and after fixing, the cells were washed 6 times in 1X PBST (1X PBS, 0.1% Tween 20), and only following PFA fixation, cells were permeabilized with 0.1% Triton X-100. After fixing, cells were blocked in 1X PBST containing 5% horse serum (ThermoFisher, 16050) for 30 min, and finally washed 6 times in 1X PBST. Cells were incubated in a wet chamber with a primary antibody solution 1:100 diluted in 5% horse serum in 1X PBST, overnight at 4°C. Cells were then washed 6 times in 1X PBST and incubated in the secondary antibody solution with dilution 1:2000 in the dark, for 2 h at room temperature. After incubation, cells were washed 6 times with 1X PBST, incubated in 10 μM Hoechst 33342 solution (Sigma-Aldrich, 14533) for 1 min, washed again 6 times, and embedded in ProLong Gold antifade reagent (ThermoFisher, P36931). Confocal imaging was performed using a Leica STELLARIS 8 FALCON system. Images were analyzed, quantified, and prepared with Fiji (v.2.3.0) and Imaris (v.10.0.0, Bitplane, Oxford Instruments), employing intensity thresholding, size exclusion, and noise filtering, based on signal intensities of the control. P62/SQSTM1 and WIPI2 foci were detected using the Spots tool based on the most intense fluorescent regions (growing spots with a minimal radius of 0.2 μm, no vertical correction applied). Abnormal or dead cells were manually excluded. Nuclei were detected using the Surface tool, and nuclei close to the edges were excluded. P62/SQSTM1 and WIPI2 foci belonging to excluded cells were then manually excluded before automated counting was applied. A minimum of 100 cells were analyzed per condition and replicate. Spots and surfaces were detected automatically (“per batch”) using the same parameters throughout each experiment to avoid any user-induced bias.

BAG2 and CLCC1 colocalization was determined using the Imaris Coloc extension for intensity-based colocalization (Imaris v.10.0.0, Bitplane, Oxford Instruments). 20 cells were analyzed per condition and biological replicated and significance was determined with a student’s t-test using Excel (Microsoft, v. 16.73). Single cell images were generated using Fiji (v.2.9.0) and Adobe Photoshop (v.22.5.1) softwares. Background subtraction, gaussian filter and brightness/contrast adjustment were applied identically to each picture displayed on a same panel.

To determine ER localization of AMBRA1, ERGFP expressing HeLa cells were examined with an LSM 900, Airyscan SR Zeiss confocal microscopy. The area of colocalizing ERGFP and AMBRA1 (Alexa647) signals was measured using ZEN 3.0 Blue edition software and expressed as μm^2^ per cell. A minimum of 25 cells/sample was analyzed and the statistical analysis was performed using ANOVA 2-way test for repeated samples by using Graphpad Prism, p values of less than 0.05 were considered significant.

### Cell lysis

For immunoblotting analyses of whole cell lysates and Halo-LC3 tag assays, cells from 1X10 cm dish were lysed in modified RIPA with SDS (50 mM Tris, 150 mM NaCl, 2% SDS, 0.5% sodium deoxycholate, 1% Triton X-100, EDTA-free 1x protease inhibitors (Roche, 11-697-498-001), 1x PhosSTOP (Roche, 04-906-837-001, 1:5000 Benzonase), pH-7.5). For HA-based affinity purifications of ULK1 complex members (ULK1, ATG101, ATG13 and FIP200), SILAC labelled cells of 2×15 cm dishes for each label were lysed in normal lysis buffer (50 mM Tris, 150mM NaCl, 1 mM EDTA, 0.1% SDS, 0.5% sodium deoxycholate, 1% Triton X-100, EDTA-free 1x protease inhibitors (Roche, 11-697-498-001), pH-7.5). For HA-based affinity purifications of SHA-BAG2 WT and phospho-variants, 2×15 cm dish of cells per replicate were lysed in BAG2 IP buffer (50 mM Tris, 150 mM NaCl, 1 mM EDTA, 1 mM EGTA, 0.35% Triton X-100, EDTA-free 1x protease inhibitors (Roche, 11-697-498-001), pH-7.5). Similar lysis buffer conditions were used for HA-based affinity purification of SHA-BAG2 in presence and absence with 20 μM of the ULK1 inhibitor MRT68921 (Merck, SML1644) (2 h prior HBSS treatment, total of 4 h). For endogenous AMBRA1 immunoaffinity purifications (antibody ABC131, Millipore), 2×15 cm dishes of cells per replicate were lysed in AMBRA1 IP buffer (10 mM Tris, 150 mM NaCl, 10% glycerol, 0.5% NP-40, EDTA-free 1x protease inhibitors (Roche, 11-697-498-001), pH-8). All cell lysates were vortexed 4 times for 15 sec, and lysis was done for 30 min on ice. Lysates were centrifuged at 14000 rpm for 10 min at 4°C. Protein quantification was determined by BCA assay as per manufacturer’s protocol (ThermoFisher, 23225). Proteins concentrations were adjusted with corresponding lysis buffers to yield equal protein amounts per lane in all respective experiments.

### Immunoblotting

Cell extracts were denatured in 1X Laemmli buffer (62.5 mM Tris pH 6.8, 2% SDS, 10% glycerol, 0.1 M DTT, 0.01% bromophenol blue), vortexed for 4 sec and heated at 75°C for 10min. For western blotting, 30 µg of total protein were loaded per well and separated by SDS-PAGE. Proteins were transferred to a 0.45 µm nitrocellulose membrane for proteins larger than 35 kDa and to 0.1 µm nitrocellulose or 0.2 µm PVDF membranes for proteins smaller than 35 kDa for 30 min using Trans-Blot Turbo transfer system (Bio-Rad, 1704150) or for 2.5 h using wet transfer system (TG buffer, VWR, 0307) under a constant electric current. Membranes were blocked for minimum 1 h in either 5% milk or 5% BSA in 1xTBST (10 mM Tris, 150 mM NaCl and 0.1% Tween-20). Membranes were incubated with the indicated primary antibodies overnight on a shaker at 4°C. Membranes were washed with 1x TBST four times 10 min and incubated with the appropriate peroxidase-conjugated secondary antibodies in 5% milk or 5% BSA in 1xTBST for 1 h at RT. Membranes were washed with 1xTBST four times 15 min. Blotted proteins were visualized using either SuperSignal West Femto chemiluminescent (ThermoFischer, PIER34096) or WesternBright ECL-(Advansta, K-12045-D50) reagents. Images were recorded with the Odyssey® Fc reader (LI-COR Biosciences-GmbH, Image Studio v.2.0.38) and densitometry analyses were performed using ImageJ Fiji software (Wayne Rasband, NIH).

### Whole proteome analyses

For whole proteome analyses of HeLa WT and HeLa KO cells (BAG2, ULK1, FIP200, ATG13, ATG101), 1X10 cm dish for each replicate per condition was lysed in 1% sodium deoxycholate, 50 mM Tris buffer, at pH=8. Samples were incubated with 1 mM DTT, heated for 10 min at 75°C followed by 5.5 mM IAA incubated for 10 min in dark at RT. Trypsin was added with 1:100 ratio (Trypsin:protein, w/w) and samples were left for digestion overnight at 37°C. TFA was added drop-wise to completely precipitate sodium deoxycholate and centrifuged at 14000 rpm, 23°C. Supernatants containing the peptides were STAGE tip-purified and dissolved in buffer A (0.1% formic acid in MS grade water) for LC-MS/MS measurements.

### Cell fractionation

Cells of 2×15 cm dishes for each cell line per replicate were cultured 36 h prior to indicated treatments and culture medium was changed 16 h before the treatments. Cell pellets were dissolved in homogenization (HM) buffer (0.25 M Sucrose, 1 mM EDTA, 20 mM HEPES-NaOH pH 7.4), supplemented with EDTA-free 1x protease inhibitors (Roche, 11-697-498-001). The cells were taken up in HM buffer and dounced 550 times followed by several centrifugation steps of supernatants from the preceding fractions to collect nuclear (1’000 g, 15 min, 4°C), mitochondria (3’000 g, 15 min, 4°C), and vesicular fractions (17’000 g, 15 min, 4°C). The last supernatant was collected as the cytosolic fraction. 1% of whole cell lysates were taken from respective samples prior fractionations. All pellets of individual fractions were washed twice with HM buffer. The pellets from mitochondria and vesicular fractions were resuspended in RIPA buffer (50 mM Tris, 150 mM NaCl, 1% Triton X-100, 0.5% sodium deoxycholate, pH 7.5, 2% SDS). The nuclear pellet was solved in 3 ml of S1 (0.25 M Sucrose, 10 mM MgCl2). This solution was layered over 3 ml of S2 (0.35 M Sucrose, 0.5 mM MgCl2), and passed through at 1’430 g for 5 min at 4°C. The nuclear pellet was solved in RIPA buffer as mentioned above. The total protein amount was quantified for all collected fractions by BCA Protein Assay (PierceR, ThermoFisher scientific) according to the manufacturer’s protocol. 30 µg total protein were taken per sample for each fraction and western blot analysis was performed as mentioned in the immunoblotting section.

### HA affinity purification and MS sample preparation

Transgenes encoding HA-miniTurbo tagged ULK1, ATG13, ATG101, FIP200, and HA-MT-GFP (pDS48), as negative ctrl were inducible expressed using 2 μg/ml doxycycline for 24 h in their respective CRISPR KO cells. 2x 15cm dishes of cells per SILAC label per replicate were used to perform HA-immunoprecipitation using 60 µl slurry of Anti-HA antibody coated magnetic beads (ThermoFisher, 13474229). Cells were lysed in normal lysis buffer; proteins were BCA quantified and equal amount of protein was taken for each label for respective replicates and affinity purification was performed at 4°C on a rotor for 4 h. Further, the beads were washed with lysis buffer and proteins were eluted twice with 30 µl Laemmli buffer at 75°C with 1 mM DTT. The SILAC labels of a replicate were mixed, proteins alkylated with 5.5 mM IAA and fractionated on 4-12% gradient gels. Gel pieces were cut into five fractions, proteins were in-gel digested with trypsin (Promega, V5113). Tryptic peptides were purified by STAGE tips prior to LC-MS/MS measurements. For HA-affinity purification of BAG2 WT and phospho-variants, a similar protocol was performed with few modifications: beads were washed 4 times with lysis buffer after affinity purification, treated with 8 M urea with 1 mM DTT in 50 mM Tris buffer pH 8, transferred onto a 10 kDa MW cut-off filter (Vivacon 500, 10,000 MWCO, Sartorius Cat#VN01H02) and spun at 10’000 g for 30 min. Proteins on the filter were alkylated with 5.5 mM IAA, spun at 10K g for 30 min and washed twice with 50 mM Tris buffer. Protein digestion for MS analysis was performed overnight according to the FASP protocol (Wiśniewski et al., 2009). Peptides were eluted twice with 200 µl with 50 mM Tris buffer into fresh tubes by centrifugation at 14’000 *g* for 30 min. Eluates were acidified with 5 µl formic acid to a final concentration of ∼1% and desalted by STAGE tips prior to LC-MS/MS measurements.

### Proximity labeling and streptavidin-based affinity purifications

Expressions of N/C-terminal fusions genes encoding HA-miniTurbo fused to ULK1, ATG13, ATG101 and FIP200 (i.e. gene-of-interest (GOI)), were induced by 2 μg/ml doxycycline, in heavy, medium, light SILAC labeled cells for 24 h. All the GOIs were expressed in their respective CRISPR KO cells. Heavy and medium labeled cells were expressing N-terminal and C-terminal fusion constructs of the above GOIs and for one out of three replicates the labels were swapped. Light labeled cells were always kept as negative control expressing doxycycline-inducible miniTurbo-GFP fusion construct in their respective knockout cells. All these cells were passaged 12 h before induction and 36 h prior to HBSS treatment for 90 min. In parallel, cells were incubated with 400 µM biotin (Sigma Aldrich, B4501) for 90 min along with 10 nM concanamycin A. Cells were washed with 1X PBS on ice for 5 times before harvesting by centrifugation at 3’000 rpm for 2 min and snap freezing in liquid nitrogen before storing the cell pellets at −80°C. For streptavidin-based affinity purification of respective proteins-of-interest (POI), 2x 15 cm dishes per label and replicate were lysed in RIPA buffer (50 mM Tris, 150 mM NaCl, 1 mM EDTA, 1 mM EGTA, 1% triton-X-100, 0.5% sodium deoxycholate, 0.1% SDS, EDTA-free 1X protease inhibitors (Roche, 11-697-498-001), pH-7.5, 1:5000 Benzonase and 1 mM PMSF). All the above cell lysates were vortexed 4 times for 15 sec, and lysis was done for 30 min on ice. Then, lysates were centrifuged at 14’000 rpm for 10 min at 4°C. Protein quantification was determined by BCA assay as per manufacturer’s protocol (ThermoFisher, 23225). Equal amounts of proteins were taken in their respective experiments to perform streptavidin (ThermoFischer, Pierce 20361) affinity purification as follows. 100 µl bead slurry was washed 2x with RIPA buffer (without 0.5% sodium deoxycholate, protease inhibitors, PMSF, 0.1% SDS) at RT. Lysates were incubated with beads on a rotor for 90 min. After affinity purification, beads were washed five times with RIPA buffer, three times with TAP lysis buffer (50 mM HEPES-KOH pH 8, 100 mM KCl, 10% glycerol, 2 mM EDTA, 0.1% Triton X-100) and two times with 50 mM ammonium carbonate. Elutions from the beads were performed using Laemmli buffer with 1 mM DTT and 25 mM biotin 75°C for 10 min. SILAC eluates of a respective experiment were mixed to give 150 µl final volume. Protein samples were reduced with 1 mM DTT followed by alkylation with 5.5 µM IAA and ran on 4-12% gradient gels. Gel pieces were sliced to have five fractions per replicate, in-gel digestion with trypsin (Promega) overnight. Peptides were purified via STAGE tips (C18 disc, 3M Empore, Cat#14-386-2) and dissolved in buffer A (0.1% formic acid in MS grade water) prior to LC-MS/MS analysis.

### Mass spectrometry-based proteomic analyses

The LC-MS/MS measurements were performed on a QExactive Plus or HF-X mass spectrometer (Thermo Scientific) coupled to an EasyLC 1000 nanoflow-HPLC. Peptides were separated on fused silica HPLC-column tips (I.D. 75 μm, New Objective, self-packed with reprosil-Pur 120 C18-AQ, 1.9 μm [Dr. Maisch] to a length of 20 cm) using a gradient of buffer A (0.1% formic acid in water) and buffer B (0.1% formic acid in 80% acetonitrile in water). Mass spectrometers were operated in the data-dependent mode; after each MS scan (mass range m/z = 370 – 1750; resolution: 70’000 for QE Plus and 120’000 for HF-X) a maximum of ten, or twelve MS/MS scans were performed using a normalized collision energy of 25%, a target value of 1’000 (QE Plus)/4500 (HF-X) and a resolution of 17’500 for QE Plus and 30’000 for HF-X. MaxQuant software (version 1.6.2.10) (Cox and Mann, 2008) was used for analyzing the MS raw files for peak detection, peptide quantification and identification using full length Uniprot human databases (version April 2016/ Jan 2022) and common contaminants. Carbamidomethylcysteine was used as fixed modification and N-terminal acetylation, oxidation of methionine, and in case of phosphorylation site analyses phosphorylation of serine, threonine or tyrosine were set as variable modifications. The MS/MS tolerance was set to 20 ppm and four missed cleavages were allowed for Trypsin/P as enzyme specificity. Based on a forward-reverse database, protein and peptide FDR was set to 0.01, minimum peptide length was set to seven, and at least one unique peptide had to be identified. The match-between run option was set to 0.7 minutes. For whole proteome analyses on the HF-X, the mass spectrometer was operated in the data-independent mode. Briefly, after each survey scan (mass range m/z = 350 – 1,200; resolution: 120,000) 28 DIA scans with an isolation width of 31.4 m/z were performed covering a total range of precursors from 350-1,200 m/z. AGC target value was set to 3 x 10^6^, resolution to 30,000 and normalized collision energy to 27%. Data were analyzed using Spectronaut software version 15.7 (Biognosys) with standard settings (without imputation) in directDIA mode using reference proteome of Human (UniProt, 2022, full length) and common contaminants. MaxQuant and DIA results were analyzed using Perseus software (version 1.6.2.3) (Tyanova et al., 2016).

The MS proteomics data have been deposited to the ProteomeXchange Consortium via the PRIDE partner repository (Perez-Riverol et al., 2022).

### MS quantification and statistical analysis

To determine significantly enriched interactors or neighbors of the ULK1 holo-complex by AP-MS or PL-MS, respectively, following criteria were used: a minimum of two valid values per 24 single AP or PL experiments were required. Significantly enriched proteins were determined by outlier tests (Significance A, p<0.05 for AP and p<0.05 BH-corrected for PL-MS, respectively). Proteins had to be significantly enriched in minimal two biological replicates of one specific bait protein.

For the BAG2 interactome analysis, label free quantification measurements were obtained for all samples. All POI APs were bait normalized and the vector control or anti-HA beads control were median normalized. Missing values in control experiments were imputed by normal distributions with a width of 0.3 and a down shift of 1.8. Proteins with FDR corrected q<0.05 values were considered significantly enriched in BAG2 WT and phospho-variants comparing against vector control or anti-HA beads control. These significantly enriched proteins were compared in stimulus-dependent APs (nutrient-rich and starvation conditions): BAG2 WT and phospho-variants BAG2 S31A and BAG2 S31E. Proteins significantly enriched with q<0.05 and p<0.05 were highlighted in respective figures. Perseus software (Version 1.6.2.3) was used for all statistical analysis.

### Halo-LC3 assays

The indicated Halo-rLC3B transgenic cells (HeLa WT cells, BAG2 KO cells and BAG2 KD cells with respective scrambled control) were induced for Halo-rLC3B expression for 24 h in 2 μg/ml doxycycline, pulsed for 20 min with 100 nM TMR ligand (Promega, G8251), washed twice in 1x DPBS, and compared in nutrient-rich and starvation conditions for 3 h at 37°C. Western blot analysis was performed for all samples (see above). Experiments were performed in triplicates and autophagy flux was calculated by taking a ratio of densitometric measurements of Halo-only bands in the presence/absence of TMR ligand in the respective cells.

### Yeast two-hybrid (Y2H) interaction analysis

For Y2H interaction assays, plasmids expressing bait proteins, fused to the Gal4 DNA-binding domain (G4BD), and prey proteins, fused to the Gal4 activation domain (G4AD), were co-transformed into reporter strain PJ69-4A (Table S2B) (James et al., 1996). Y2H interactions were documented by spotting representative transformants in 10-fold serial dilution steps onto SC-Leu-Trp (-LT), SC-Leu-Trp-His (-HLT; *HIS3* reporter), and SC-Leu-Trp-Ade (-ALT; *ADE2* reporter) plates, which were incubated for 3 d at 30°C. Growth on SC-Leu-Trp-His plates is indicative of a weak/moderate interaction, whereas only relatively strong interactions permit growth on SC-Leu-Trp-Ade plates. Y2H plasmids were constructed by standard cloning procedures using *Escherichia coli* DH5α for cloning and plasmid propagation. All cloned DNA fragments generated by PCR amplification were verified by sequencing. More information on the plasmids, which are listed in Supplemental table S2E, and for yeast strains in Supplemental table S2D, are available upon request.

## Acknowledgement

This work was supported by the Canton and the University of Fribourg as part of the SKINTEGRITY.CH collaborative research project and by the Swiss National Science Foundation (grants 310030_212187, 310030_184781 to JD, grant 310030_192614 to WJK). This work has been also supported by grants from the Italian Ministry of Health (Ricerca Corrente e Finalizzata, GR-2019-12369231, PNRR-MAD-2022-12375755, MA and GMF), and from CAPES, Coordenação de Aperfeiçoamento de Pessoal de Nível Superior – Brasil (Código de Financiamento 001) for scholarship to LODM. We acknowledge co-funding from Next Generation EU through the Italian Ministry of University and Research, in the context of the National Recovery and Resilience Plan, Investment PE8 – Project Age-It: “Ageing Well in an Ageing Society” to GMF, co-financed by the Next Generation EU (DM 1557 11.10.2022) CUP B53C22004090006. We would like to thank Prof Mario Tschan, Dr Anna Bill and Dr. Jun Xu for the generous support with FACS analysis. We thank Michael Stumpe for mass spectrometry support.

