## Supplemental Figures for "The ULK1 effector BAG2 regulates autophagy initiation by modulating AMBRA1 localization"

**Supplemental Figure S1:** Streptavidin enrichment of miniTurbo-based biotinylation reactions of ULK1 complex members.
**Supplemental Figure S2:** Deep ULK1 complex interactome.
**Supplemental Figure S3:** Assessment of protein-protein interactions by yeast two-hybrid (Y2H) assays.
**Supplemental Figure S4:** Perturbed autophagy flux upon loss of BAG2.
**Supplemental Figure S5:** Perturbed autophagosomal-lysosomal targeting upon chronic loss of BAG2.
**Supplemental Figure S6:** Increased colocalization of BAG2 and SEC16A in starvation conditions.
**Supplemental Figure S7:** Colocalization of CLCC1 with organellar markers.
**Supplemental Figure S8:** Increased number of WIPI2 puncta upon acute depletion of BAG2.
**Supplemental Figure S9:** Increased colocalization of $BAG2^{S31E}$ and CLCC1 in starvation conditions.

**Supplemental References**

**Suppl Figure S1**

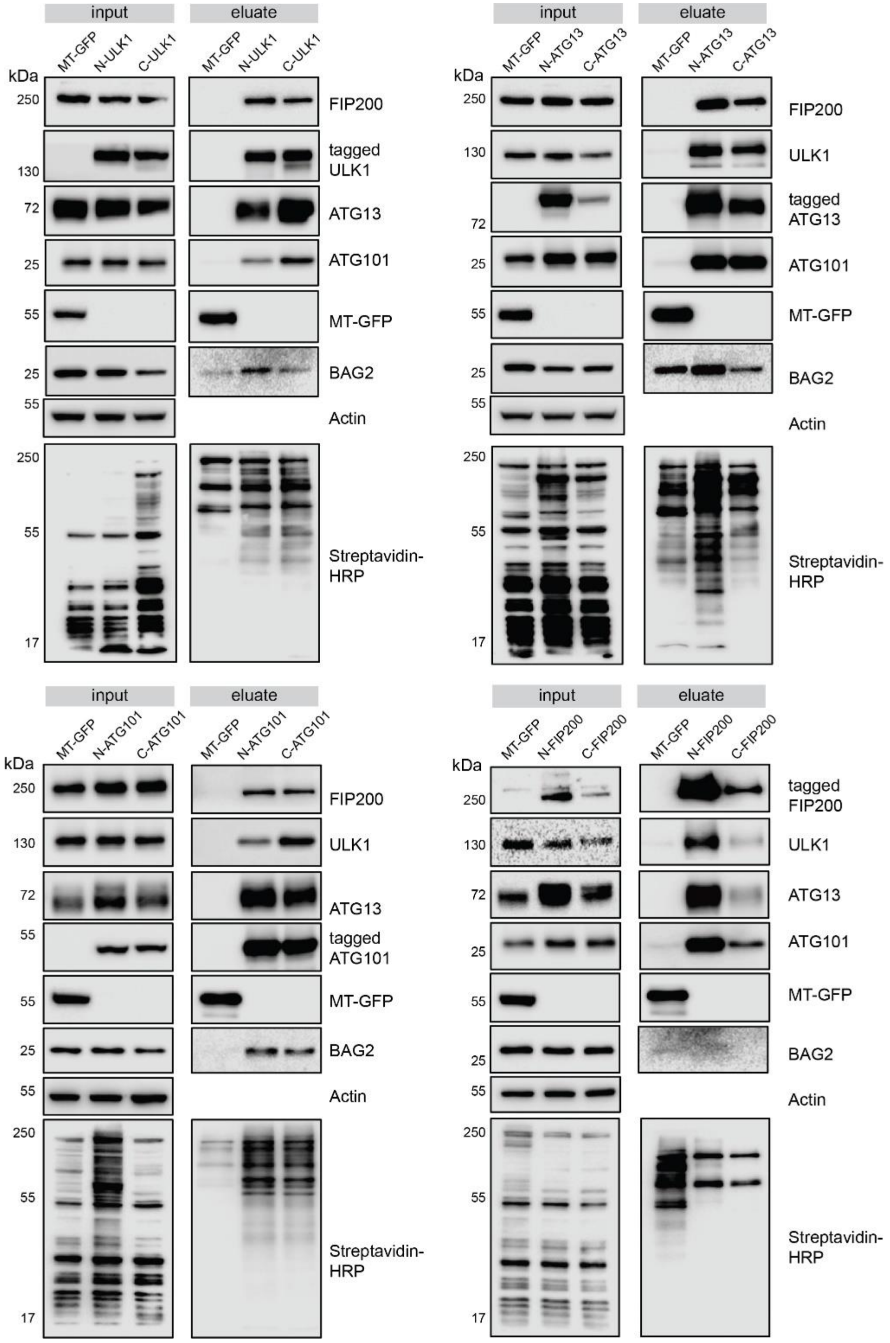


**Supplemental Figure S1: Streptavidin enrichment of miniTurbo-based biotinylation reactions of ULK1 complex members.** CRISPR KO HeLa cells of ULK1 complex members were used to express indicated N-/C-terminal miniTurbo fusion constructs. Streptavidin-based APs were performed followed by western blot analysis. Input and eluates are shown in respective figures and Streptavidin-HRP was used to detect biotinylation reactions. MT-GFP was used as negative control in respective CRISPR KO cell lines along with the indicated bait proteins.

**Suppl Figure S2**

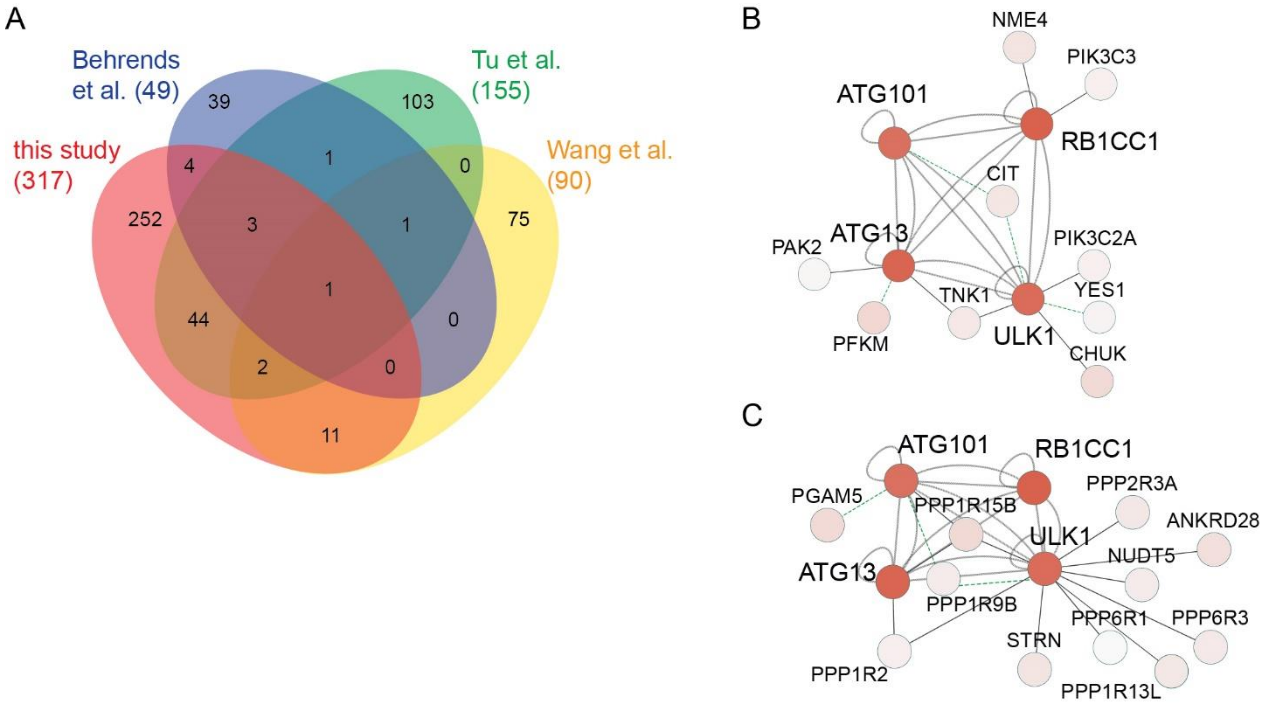


**Supplemental Figure S2: Deep ULK1 complex interactome. (A)** Comparison of the current ULK1 complex deep interactome with existing studies. The overlap of the current studies with published data is larger than the overlap between respective data sets (Behrends et al., 2010; Tu et al., 2021; Wang et al., 2019). Numbers in brackets indicate total numbers of hits of respective studies. **(B)** Protein- and lipid-kinases identified as ULK1 complex interactors in the current study (see Figure 2). **(C)** Phosphatase subunits identified as ULK1 complex interactors in the current study (see Figure 2).

**Suppl Figure S3**

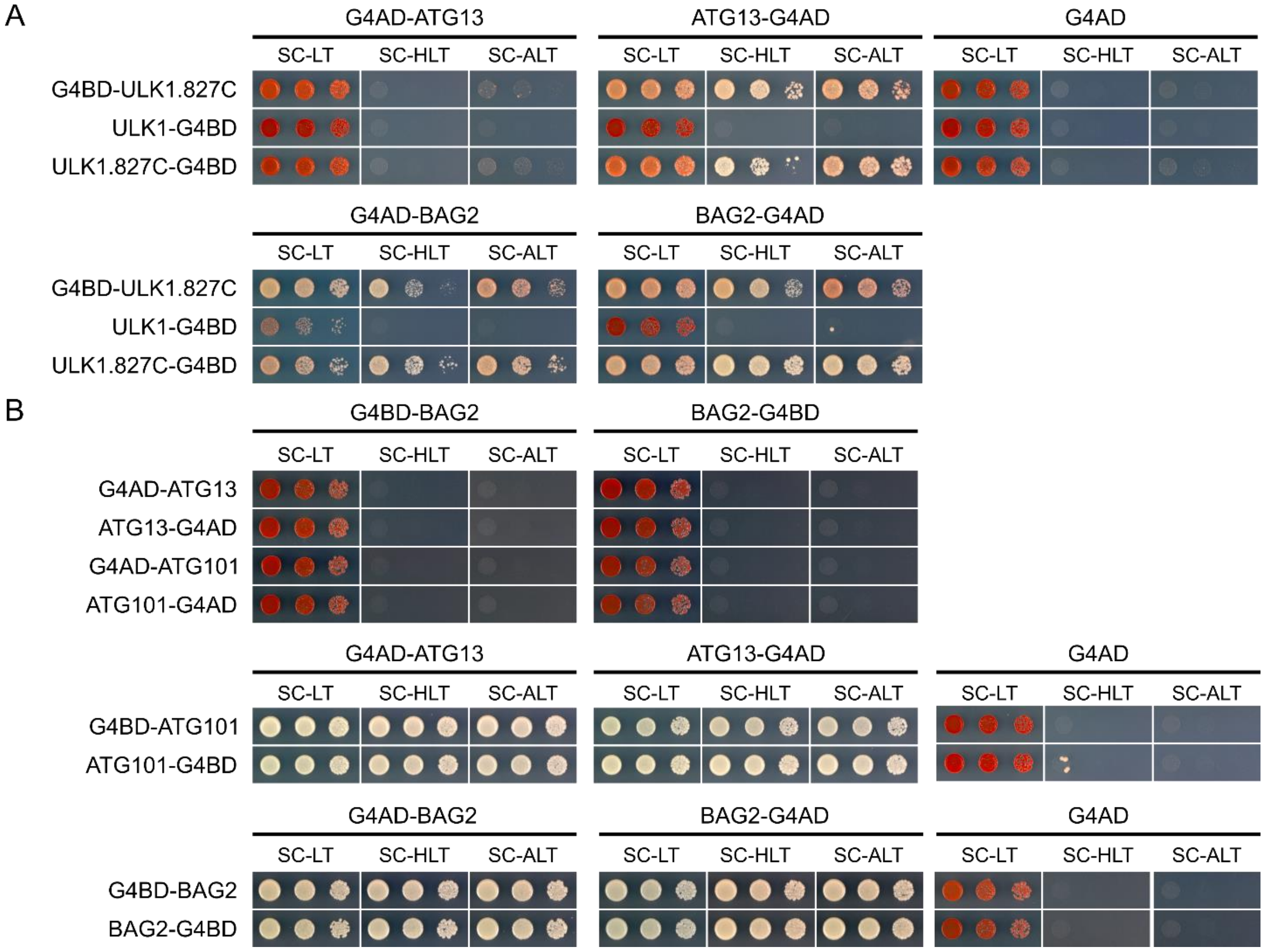


**Supplemental Figure S3: Assessment of protein-protein interactions by yeast two-hybrid (Y2H) assays.** The Y2H reporter strain PJ69-4A was transformed with two plasmids expressing the human proteins of interest fused to the Gal4 activation domain (G4AD) or Gal4 DNA-binding domain (G4BD) at either their N- or C-terminal end. Transformed cells were spotted in 10-fold dilutions series onto SC-Leu-Trp (SC-LT), SC-His-Leu-Trp (SC-HLT), SC-Ade-Leu-Trp (SC-ALT) plates, which were incubated for 3 d at 30°C. **(A)** ATG13 (positive control) and BAG2 interact with the C-terminal domain of ULK1 (abbreviated as ULK1.827C); however, no interaction is observed with full-length ULK1. **(B)** BAG2 neither directly interacts with ATG13 nor ATG101, while the known/established ATG13-ATG101 and BAG2-BAG2 interactions could be recapitulated by the Y2H assays. Note that none of the tested fusion proteins exhibit self-activation of the reporter genes, as indicated by the absence of growth when these are co-expressed with the non-fused G4AD or G4BD.

**Suppl Figure S4**

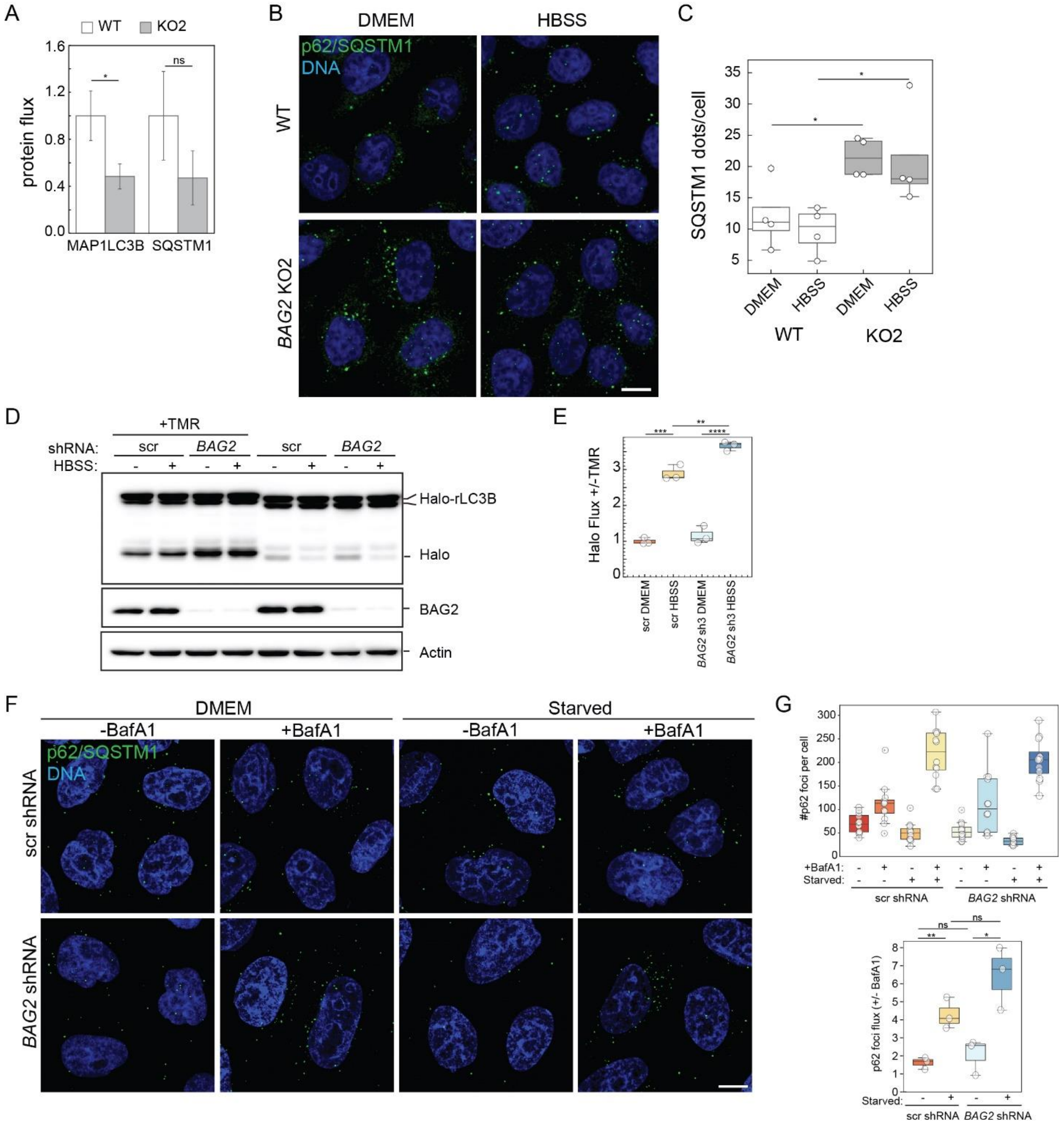


**Supplemental Figure S4: Perturbed autophagy flux upon loss of BAG2. (A)** KO of BAG2 reduces protein turnover. Quantitative MS data of n=3 biological replicates. Cells were starved for 2 h with or without the presence of 10 nM ConA. Protein fluxes represent the abundance ratios of ConA compared to non-ConA samples. Error bars indicate SD. *: p<0.05, LIMMA. **(B-C)** SQSTM1/p62 dot formation analyzed by IF. Fluorescent micrographs showing p62/SQSTM1 foci formation in *BAG2* WT and KO HeLa cells in untreated (DMEM) or starved (HBSS, 3h) conditions (B). Scale bar= 10 μm. One representative image out of n=4 biological replicates is shown. (C) Quantification of p62/SQSTM1 foci count per cell as shown in (B). White dots represent the average foci count per cell and per biological replicate (=4). A minimum of 40 cells were counted per replicate. An unpaired two-tailed Student's t-test was used to compare values. *: $p \leq 0.05$. **(D)** Halo-LC3 assay using WT HeLa cells transgenic for an inducible Halo-LC3 construct, transduced with either *scrambled*- or *BAG2* shRNA-expressing lentiviruses, then pulsed for 30 min with 100 nM TMR and either incubated in full growth medium (DMEM) or starvation medium (HBSS) for 3 h. One representative assay out of n=3 biological replicates is shown. An increase in molecular weight is observed for the Halo-LC3 and Halo fragments when TMR ligand is conjugated. **(E)** Quantification of the Halo autophagic flux in (D), corresponding to the ratio of Halo tag released in presence vs in absence of TMR (+/-). White dots: individual biological replicates, n=3. An unpaired two-tailed Student's t-test was used to compare values. **: $p \leq 0.01$; ***: $p \leq 0.001$; ****: $p \leq 0.0001$. **(F)** Fluorescent micrographs showing p62/SQSTM1 foci formation in WT HeLa cells transduced with either *scrambled*- or *BAG2* shRNA-expressing lentiviruses, in untreated (DMEM) or starved (HBSS, 3h) conditions supplemented or not with 2nM BafA1. Scale bar= 10 µm. One representative image out of n=3 biological replicates is shown. **(G)** Quantification of p62/SQSTM1 foci count per cell as shown in (F). White dots represent the average foci count per cell and per image. 4 images were quantified per replicate of n=3 biological replicate. An unpaired two-tailed Student's t-test was used to compare values. *: $p \leq 0.05$; **: $p \leq 0.01$.

**Suppl Figure S5**

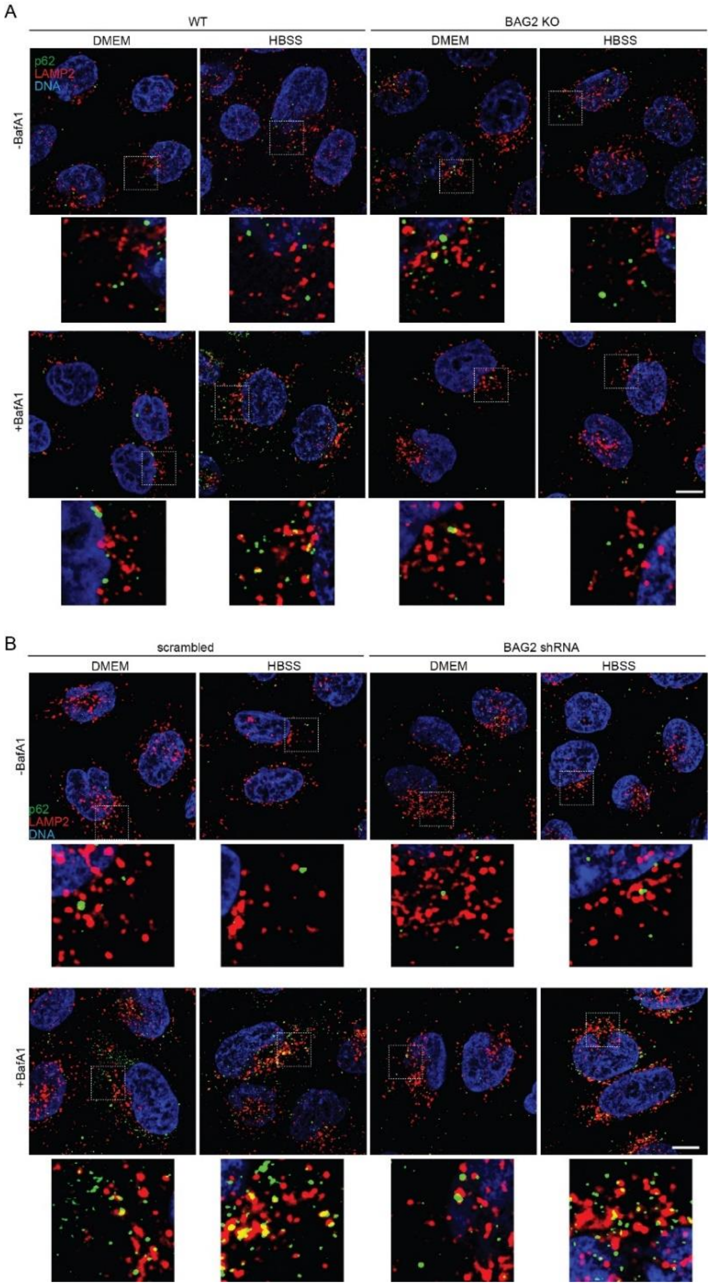


**Supplemental Figure S5: Perturbed autophagosomal-lysosomal targeting upon chronic loss of BAG2.** Fluorescent micrographs labeled with indicated primary antibodies. In BAG2 KO cells reduced colocalization of p62 and LAMP2 in starvation (HBSS) and BafA1 treatment is observed compared to WT cells (A). In WT HeLa cells transduced with either *scrambled*- or *BAG2* shRNA-expressing lentiviruses, no difference is observed (B). Untreated (DMEM) or starved (HBSS, 3 h) conditions supplemented or not with 2 nM BafA1 are analyzed. Scale bar= 10 μm. One representative image out of n=3 biological replicates is shown. Respective quantification is shown in main Figure 3J-K.

**Suppl Figure S6**

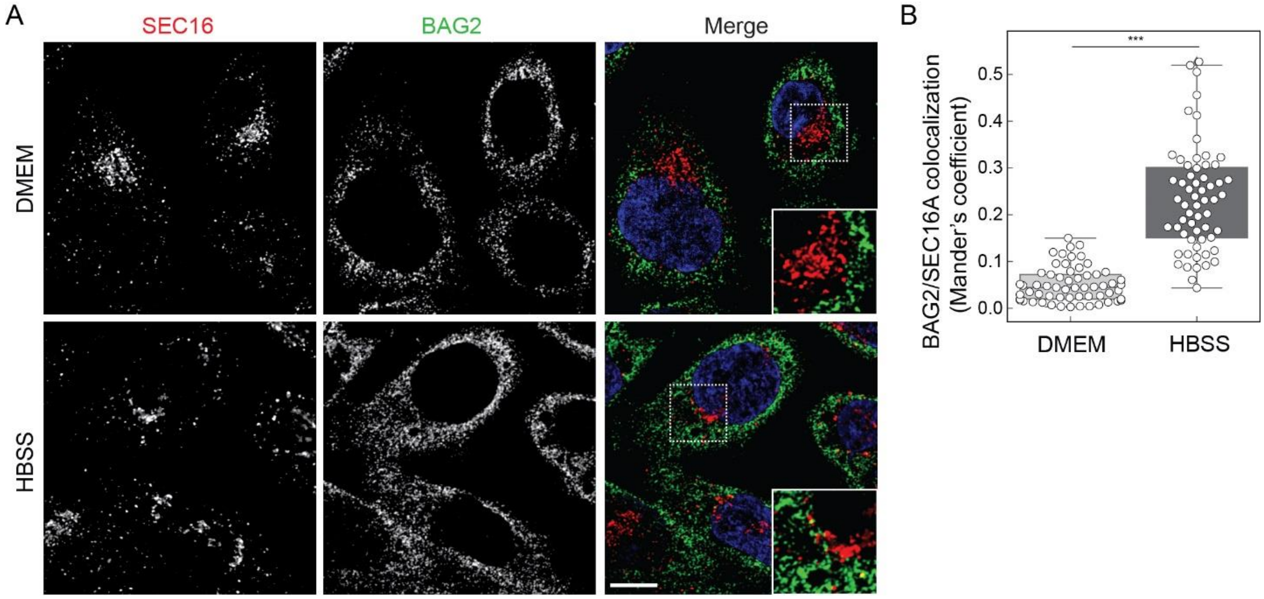


**Supplemental Figure S6: Increased colocalization of BAG2 and SEC16A in starvation conditions. (A)** Fluorescent micrographs labeled with indicated primary antibodies. In starved cells (HBSS, 2 h) the BAG2/SEC16A colocalization increases compared to fed cells (DMEM), indicating increased BAG2 localization at the ER membrane. **(B)** Quantification of (A). A total of 60 cells per condition were measured for BAG2/SEC16A colocalization in n=3 biological replicates and are represented by white dots; representative pictures are shown. An unpaired student's t-test was used to determine the significant differences. ***: $p<0.001$. Scale bar= 10 μm.

**Suppl Figure S7**

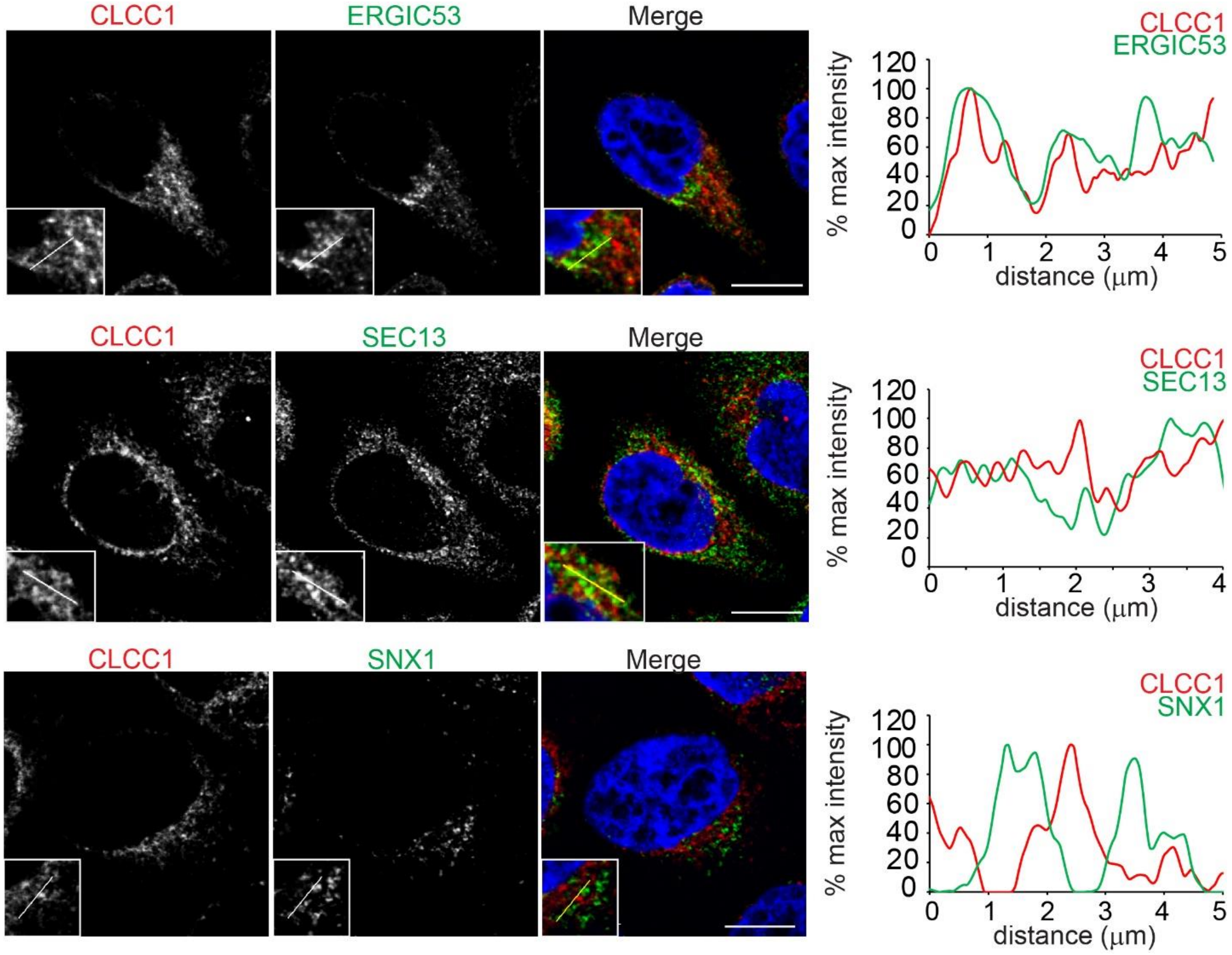


**Supplemental Figure S7: Colocalization of CLCC1 with organellar markers.** CLCC1 partially colocalizes with ERGIC53 and SEC13, but not with SNX1. Left panels: fluorescent micrographs from HeLa WT cells in fed conditions, labeled using the indicated primary antibodies. Yellow lines: line drawings used to generate plot profiles. Right panels: plot profiles corresponding to the yellow lines. The signal of each fluorophore was normalized to their maximum intensity to obtain comparable profiles between antibodies. Scale bar= 10 μm.

**Suppl Figure S8**

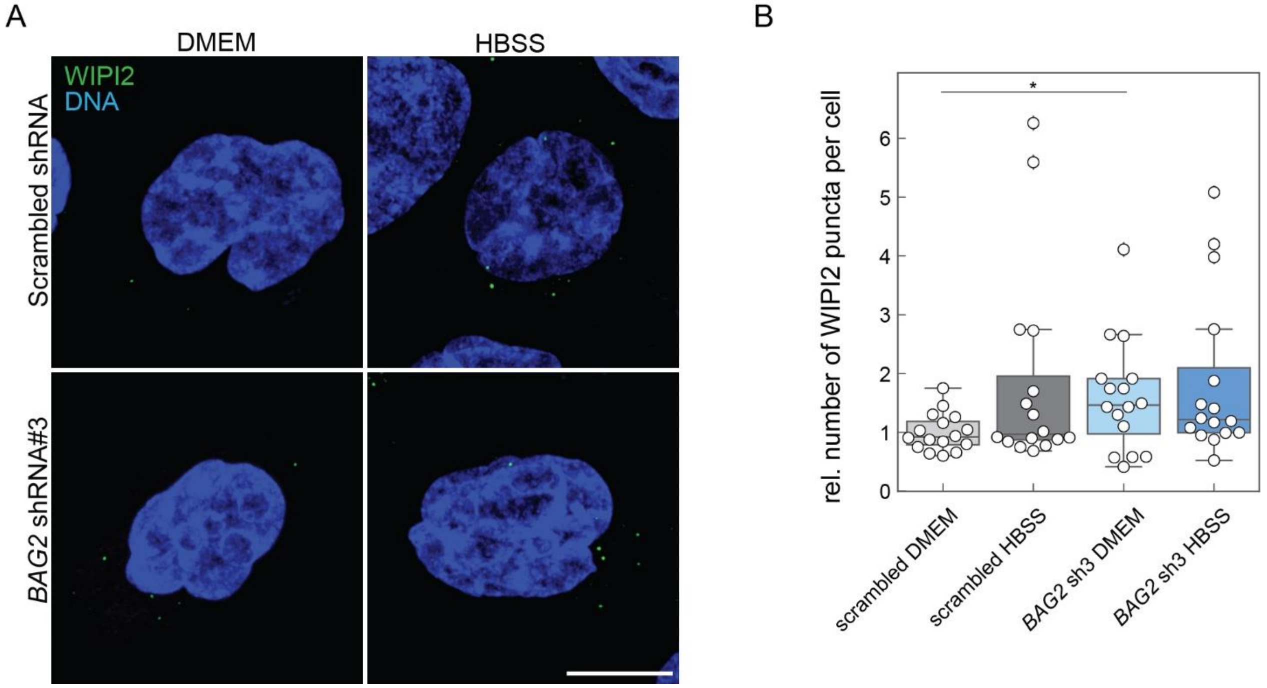


**Supplemental Figure S8: Increased number of WIPI2 puncta upon acute depletion of BAG2. (A-B)** Acute loss of BAG2 leads to increased number of WIPI2 dots. (A) WIPI2 puncta analysis using IF micrographs from untreated (DMEM) or starved (HBSS, 3 h) HeLa WT treated with a scrambled control shRNA and an anti-BAG2 shRNA. Representative pictures of n=4 biological replicates are shown. Scale bar= 10 μm. (B) Puncta quantification of (A). Each white dot represents the quantified average WIPI2 dots per cell per image (8 to 19 cells per image), with a total of 16 images (4 images per condition), per replicate of n=4 biological replicates. An unpaired student's t-test was used to determine the significant differences. *: $p<0.05$. Scale bar = 10 μm

**Suppl Figure S9**

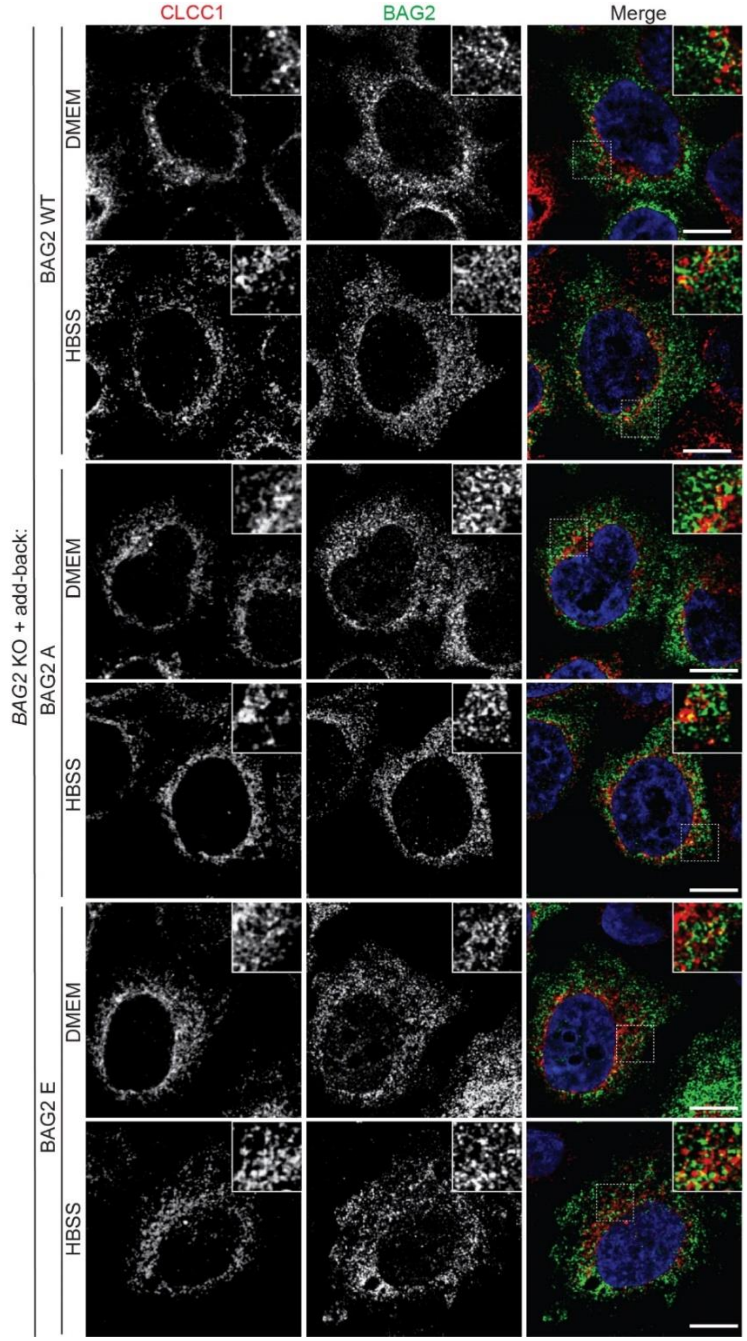


**Supplemental Figure S9: Increased colocalization of BAG2$^{S31E}$ and CLCC1 in starvation conditions.** Images corresponding to Figure 6F quantifications. HeLa *BAG2* KO cells transduced with either WT, S31A ("A") or S31E ("E") BAG2 variant add-back constructs were either left in full growth medium (DMEM) or starved for 3 h (HBSS) before being processed for immunofluorescence with the indicated antibodies. One representative set of pictures out of n=3 biological replicates is displayed. Scale bar= 10 μm.
