## Supplemental Table S2 for "The ULK1 effector BAG2 regulates autophagy initiation by modulating AMBRA1 localization"

| Table S2A: List of primers for PCR, and cDNA regarding cloning | | |
| --- | --- | --- |
|  | **PCR primer numbers** | **Primer sequences** |
| 1 | ONDS115 | CAGATCGCCTGGAGAATTGGCTAGCCATATGGCCACCATGTACCCATACGATGTTCCAGATTACGCTATCCCGCTGCTGAACGCTAAACAG |
| 2 | ONDS116 | CAGGCCCAGCAGGGGGTTGGGGATGGGCTTGCCCTTTTCGGCAGACCGCAGACTGATTTC |
| 3 | ONDS117 | CCCATCCCCAACCCCCTGCTGGGCCTGGACAGCACCGAGCCCGGCCGCGG |
| 4 | ONDS118 | CATATCCAGTCACTATGGTCGACCTGCAGGGATCCTTATCAGGCACAGATGCCAGTCAGC |
| 5 | ONDS119 | CCCATCCCCAACCCCCTGCTGGGCCTGGACAGCACCGAAACTGATCTCAATTCCCAGGACAGAAAGG |
| 6 | ONDS120 | CATATCCAGTCACTATGGTCGACCTGCAGGGATCCTTACTATTGCAGGGTTTCCACAAAGGCATC |
| 7 | ONDS121 | CCCATCCCCAACCCCCTGCTGGGCCTGGACAGCACCAAGTTATATGTATTTCTGGTTAACACTGGAACTACTC |
| 8 | ONDS122 | CATATCCAGTCACTATGGTCGACCTGCAGGGATCCCTATTATACTTTCTTATTCCATGATACGGCTTTCAC |
| 10 | ONDS132 | CAGGCCCAGCAGGGGGTTGGGGATGGGCTTGCCCATGGATCCAGCCCAGTTCTCCAATTC |
| 11 | ONDS133 | CCCATCCCCAACCCCCTGCTGGGCCTGGACAGCACCATCCCGCTGCTGAACGCTAAACAG |
| 12 | ONDS139 | CCCATCCCCAACCCCCTGCTGGGCCTGGACAGCACCGTGAGCAAGGGCGAGGAG |
| 13 | ONDS140 | CATATCCAGTCACTATGGTCGACCTGCAGGGATCCTAACTACTTGTACAGCTCGTCCATGCC |
| 14 | ONDS163 | CAGATCGCCTGGAGAATTGGCTAGCATATGGCCACCATGTACCCATAC |
| 15 | ONDS164 | TCCAGTCACTATGGTCGACCTGCAGCTCGAGTTATCAGGCACAGATGCCAGTC |
| 16 | ONDS188 | CCGTCAGATCGCCTGGAGAATTGGCTAGCCATATGGCCACCATGAAGTTATATGTATTTCTGGTTAACACTGGAACTACTC |
| 17 | ONDS189 | CCAGGCCCAGCAGGGGGTTGGGGATGGGCTTGCCTACTTTCTTATTCCATGATACGGCTTTCAC |
| 18 | ONDS190 | CCGTCAGATCGCCTGGAGAATTGGCTAGCCATATGGCCACCATGGAGCCCGGCCG |
| 19 | ONDS191 | CAGGCCCAGCAGGGGGTTGGGGATGGGCTTGCCGGCACAGATGCCAGTCAGCAGC |
| 20 | ONDS225 | CAGATCGCCTGGAGAATTGGCTAGCCATATGGCCACCATGGAAACTGATCTCAATTCCCAGGACAGAAAGG |
| 21 | ONDS226 | CAGGCCCAGCAGGGGGTTGGGGATGGGCTTGCCTTGCAGGGTTTCCACAAAGGCATC |
| 22 | ONDS227 | ATGAACTGTCGCTCGGAG |
| 23 | ONDS228 | TCAGAGGGCAAGGGTGTC |
| 24 | ONDS229 | CAGATCGCCTGGAGAATTGGCTAGCCATATGGCCACCATGAACTGTCGCTCGGAGG |
| 25 | ONDS230 | GGCCCAGCAGGGGGTTGGGGATGGGCTTGCCGAGGGCAAGGGTGTCTTTGATG |
| 26 | ONDS231 | ATCCCCAACCCCCTGCTGGGCCTGGACAGCACCAACTGTCGCTCGGAGG |
| 27 | ONDS232 | TCCAGTCACTATGGTCGACCTGCAGGGATCCTCAGAGGGCAAGGGTGTC |
| 28 | ONDS309 | ATGGCTCAGGCGAAGATC |
| 29 | ONDS310 | CTAATTGAATCTGCTTTCAGCATTTTG |
| 30 | ONDS318 | CGATGAAAATTTATATTTTCAAGGTATGGCTCAGGCGAAGATC |
| 31 | ONDS319 | CATATCCAGTCACTATGGTCGACCTGCAGCTAATTGAATCTGCTTTCAGCATTTTG |
| 32 | ONDS345 | GTCAGATCGCCTGGAGAATTGG |
| 33 | ONDS346 | CTGCTGGAGGCTCTGGACCAG |
| 34 | ONDS347 | CTGGTCCAGAGCCTCCAGCAG |
| 35 | ONDS348 | CATATCCAGTCACTATGGTCGACC |
| 36 | ONDS349 | CTGCTGGAGGAACTGGACCAG |
| 37 | ONDS350 | CTGGTCCAGTTCCTCCAGCAG |
| 38 | ONDS379 | CAATCCATAGTAGCTGGCTGTGCTC |
| 39 | ONDS380 | GAGCACAGCCAGCTACTATGGATTG |
| 40 | ONDS107 | ACCTTGAAAATATAAATTTTCATCGGCCGCTTTTTCGAACTGC |
| 41 | ONDS111 | CAGATCGCCTGGAGAATTGGCTAGCGAATTCGCCACCATGTACCCATACGATGTTCCTGACTATGCCG |
| 42 | OSKP286 | AAAGTGGTTTAGTAATGAACCGGTAATTCTACCGGGTAGGGGAG |
| 43 | OSKP287 | GACTTTGCTCTTGTCCAGTCTAGACATTGGACCAGGGTTTTCTTCAACATCACCACAAGTGAGGAGAGAACCTCTACCTTCTTCCTTTGCCCTCGGACGAG |
| 44 | OSKP298 | ATTGGCTAGCGCCACCATGGAAATCGGTACTGGCTTTCCATTC |
| 45 | OSKP299 | ACCCGAATTCCCGGATCCCGAGCCTGAACCGCCGGAAATCTCGAGCGTCG |
| TableS2B: List of sequencing primers for all cloning | | |
|  | **Primer number** | **Primer sequences** |
| 1 | ONDS11 | GAGCCCGGCCGCG |
| 2 | ONDS12 | CCATCCCCCGGGAGACC |
| 3 | ONDS13 | GGAACCATCCCTGAGCGG |
| 4 | ONDS14 | GCGGGCCCCACTGG |
| 5 | ONDS16 | GTAGGCGTGTACGGTGGGAG |
| 6 | ONDS123 | GCCCTGCTGCTGGAAC |
| 7 | ONDS124 | TCAGAAAAAGCTGAGACGAAAAGATC |
| 8 | ONDS125 | GTCAAAGATGGAAAGAGATTATATGAAGC |
| 9 | ONDS126 | CATTAAGGAAGACCTTTGCCAC |
| 10 | ONDS127 | GTAAACTCAACCAAAAGATTCAGGATAATAATG |
| 11 | ONDS128 | GATTTTGCATAAAAAACAGACTACATAATACTG |
| 12 | ONDS198 | CCCAGGTGAATCAGATAGATCC |
| 13 | ONDS256 | TTGCAGGGTTTCCACAAAGGC |
| 14 | ONDS317 | GTTTTAAAATGGACTATCATATGCTTACCG |
| 15 | ONDS394 | CCCAGAATCTGTTTAGCGTTC |
| 16 | OSKP300 | GCGTGTACGGTGGGAGGC |
| 17 | OSKP301 | TGCTAAAGCGCATGCTCCAG |

| TableS2C: List of CRISPR sgRNA sequences and shRNA sequences | | | |
| --- | --- | --- | --- |
|  | **Primer number** |  | **sgRNA Sequence** |
| 1 | CRONDS1 | ULK1_1 | GCTACGAGAAGAACAAGACGT |
| 2 | CRONDS2 | ULK1_2 | GCAGGTTTCGCTCAATGCGC |
| 3 | CRONDS3 | ATG13_1 | ACTTGGCATTCATGTCTACCA |
| 4 | CRONDS4 | ATG13_2 | TCTTGAGAGTCTTCTACAGAC |
| 5 | CRONDS5 | ATG13_3 | GAATGGACACATTACCTTGA (Ge et al., 2017) |
| 6 | CRONDS6 | FIP200 | GTAGTTTTAGGAATAGCAGG |
| 7 | CRONDS7 | ATG101_1 | ACCAGAAGAAGAAGTCTCGCTGG |
| 8 | CRONDS8 | ATG101_2 | GTTATCCACCTCCGACTGTGTGG |
| 9 | CRONDS9 | BAG2_1 | GATCAACGCTAAAGCCAACGAGG |
| 10 | CRONDS10 | BAG2_2 | CACCTCATCAATAATCCTTGTGG |
| 11 | TRC0000033591 | shRNA-1 (BAG2) | CCGGGATCAGAAGTTTCAATCCATACTCGAGTATGGA  TTGAAACTTCTGATCTTTTTG |

**Table S2D: Yeast strain table**

| **Name** | **Mating type** | **Genotype** | **Source** |
| --- | --- | --- | --- |
| PJ69-4A | *MAT***a** | *trp1-901 leu2-3,112 ura3-52 his3-200 gal4Δ gal80Δ LYS2::GAL1-HIS3 GAL2-ADE2 met2::GAL7-lacZ* | James et al., 1996; PMID: 8978031 |

**Table S2E: Yeast Plasmid table**

| **ID** | **Name** | **Relevant information** | **Source** | **Figure** |
| --- | --- | --- | --- | --- |
| pDK2966 | pGAG4ADC111 | CEN, LEU2, PADH1, TADH1 C-terminal (GA)5-G4AD | Lab plasmid | S3B |
| pDK7500 | pG4ADHAN181 | 2µ, LEU2, PADH1, TADH1  N-terminal G4AD-HA | Lab plasmid | S3A |
| pDK10406 | pG4BDN22-HsATG101 | CEN, TRP1, PADH1, TADH1  N-terminal-G4BD | This study | S3B |
| pDK10407 | pG4BDN22-HsBAG2 | CEN, TRP1, PADH1, TADH1  N-terminal-G4BD | This study | S3B |
| pDK10410 | pGAG4BDC22-HsATG101 | CEN, TRP1, PADH1, TADH1  C-terminal (GA)5-G4BD | This study | S3B |
| pDK10411 | pGAG4BDC22-HsBAG2 | CEN, TRP1, PADH1, TADH1  C-terminal (GA)5-G4BD | This study | S3B |
| pDK10413 | pG4ADHAN111-HsATG13 | CEN, LEU2, PADH1, TADH1  N-terminal G4AD-HA | This study | S3A, S3B |
| pDK10414 | pG4ADHAN111-HsATG101 | CEN, LEU2, PADH1, TADH1  N-terminal G4AD-HA | This study | S3B |
| pDK10415 | pG4ADHAN111-HsBAG2 | CEN, LEU2, PADH1, TADH1  N-terminal G4AD-HA | This study | S3B |
| pDK10417 | pGAG4ADC111-HsATG13 | CEN, LEU2, PADH1, TADH1  C-terminal (GA)5-G4AD | This study | S3A, S3B |
| pDK10418 | pGAG4ADC111-HsATG101 | CEN, LEU2, PADH1, TADH1  C-terminal (GA)5-G4AD | This study | S3B |
| pDK10419 | pGAG4ADC111-HsBAG2 | CEN, LEU2, PADH1, TADH1  C-terminal (GA)5-G4AD | This study | S3B |
| pDK10801 | pG4BDN112-HsULK1.827C | 2µ, TRP1, PADH1, TADH1  N-terminal-G4BD | This study | S3A |
| pDK10802 | pGAG4BDC112-HsULK1 | 2µ, TRP1, PADH1, TADH1  C-terminal (GA)5-G4BD | This study | S3A |
| pDK10803 | pGAG4BDC112-HsULK1.827C | 2µ, TRP1, PADH1, TADH1  C-terminal (GA)5-G4BD | This study | S3A |
| pDK10804 | pG4ADHAN181-HsBAG2 | 2µ, LEU2, PADH1, TADH1  N-terminal G4AD-HA | This study | S3A |
| pDK10805 | pGAG4ADC181-HsBAG2 | 2µ, LEU2, PADH1, TADH1  C-terminal (GA)5-G4AD | This study | S3A |

**S2F: Sequence of rat MAP1LC3B**

ccgtccgagaagaccttcaaacagcgccggagcttcgaacaaagagtggaagatgtccggctcatccgggagcagcaccccaccaagatcccagtgattatagagcgatacaagggtgagaagcagctgcccgtcctggacaagaccaagttccttgtacctgatcacgtgaatatgagcgaactcatcaagataattagaaggcgcctgcagctcaatgctaaccaagccttcttcctcctggtgaatgggcacagcatggtgagtgtgtccacacccatctctgaagtgtacgagagcgagagagatgaagacggcttcctgtacatggtctatgcctcccaggagacgttcgggacagcactggctgttacatacatgtcagctctgaaggcaacagcaacaggaagagagccatgcttg

Ge, L., Zhang, M., Kenny, S. J., Liu, D., Maeda, M., Saito, K., . . . Schekman, R. (2017). Remodeling of ER-exit sites initiates a membrane supply pathway for autophagosome biogenesis. *EMBO Rep, 18*(9), 1586-1603. doi:10.15252/embr.201744559
